# The functional and phenotypic diversity of single T-cell infiltrates in human colorectal cancer as correlated with clinical outcome

**DOI:** 10.1101/2020.09.27.313445

**Authors:** Kazuya Masuda, Adam Kornberg, Sijie Lin, Patricia Ho, Kerim Secener, Nathan Suek, Alyssa M. Bacarella, Matthew Ingham, Vilma Rosario, Ahmed M. Al-Masrou, Steven A. Lee-Kong, P. Ravi Kiran, Kelley S. Yan, Marlon Stoeckius, Peter Smibert, Paul E. Oberstein, Peter A. Sims, Arnold Han

**Author notes:** Correspondence: Arnold Han.

## Abstract

Although degree of T-cell infiltration in CRC was shown to correlate with a positive prognosis, the contribution of phenotypically and functionally distinct T cell subtypes within tumors remains unclear. We analyzed 37,931 single T cells with respect to transcriptome, TCR sequence and 23 cell surface proteins, from tumors and adjacent normal colon of 16 patients. Our comprehensive analysis revealed two phenotypically distinct cytotoxic T cell populations within tumors, including positively prognostic effector memory cells and non-prognostic resident memory cells. These cytotoxic T cell infiltrates transitioned from effector memory to resident memory in a stage-dependent manner. We further defined several Treg subpopulations within tumors. While Tregs overall were associated with positive clinical outcomes, CD38^+^ peripherally-derived Tregs, phenotypically related to Th17 cells, correlated with poor outcomes independent of cancer stage. Thus, our data highlight the diversity of T cells in CRC and demonstrate the prognostic significance of distinct T cell subtypes, which could inform therapeutic strategies.

## Introduction

Landmark studies performed on histological samples correlated T cell-infiltration with cancer survival in colorectal cancer (CRC), a highly prevalent cancer and leading cause of cancer deaths worldwide^1,2^. However, the success of immunotherapy in CRC has been largely limited to a small subset of CRC with high microsatellite instability (MSI-H), characterized by higher degree of mutation due to defective DNA mismatch repair and accounting for ∼15% of CRC^3-7^. Several traits of MSI-H tumors have been proposed to account for its efficacy, including increased infiltration of effector CD45RO^+^CD8^+^ T cells and T-bet^+^ lymphocytes (like T-helper 1 cells), and higher degree of PD-1-PDL1 expression^5,8,9^. However, none of these traits are unique to MSI-H CRC, which is also not universally responsive to anti-PD-1 therapy^5,10^. These observations imply the feasibility of novel T cell-directed immunotherapies to treat CRC.

In contrast to CD8^+^ T cells, the contribution of CD4^+^ T-cell infiltrates in CRC to clinical outcome, in particular regulatory T cells (Tregs) and T helper 17 (Th17) cells, is controversial^11-18^. In several cancers including breast, lung, gastric and hepatocellular carcinomas, Treg infiltration negatively correlates with patient survival^19-23^. In contrast, the presence of Tregs in CRC represents a conundrum with some groups concluding that Treg density predicts positive outcomes^24,25^, although a recent report suggested different clinical outcomes were associated with heterogenous Tregs in CRC^26^.

To probe the functional and phenotypic heterogeneity of T-cell infiltrates and its contribution to tumor growth in CRC, we performed unbiased droplet-based single-cell RNA sequencing (scRNA-seq) of T cells isolated from tumors or adjacent normal colon of 16 CRC patients^18^. We used a modified workflow for scRNA-seq incorporating oligonucleotide-tagged antibodies to assess cell surface expression levels in T cells, termed CITE-seq (cellular indexing of transcriptomes and epitopes by sequencing) and also distinguish cells from different samples for multiplexing, termed cell hashing^27,28^.

Although recent scRNA-seq technologies have enabled cell-type specific RNA profiling, they still lack the scalability of bulk RNA-seq^29^. We applied semi-quantitative gene set enrichment analysis (GSEA) to quantify specific T cell types characterized by scRNA-seq from bulk RNA-seq databases in the TCGA cohort of several types of tumors including CRC, breast cancer (BRC) and melanoma^30-33^. Collectively, our study shows the dynamic relationship between phenotypically and functionally distinct T cell subtypes in CRC, and demonstrates the prognostic significance of certain T cell subtypes, providing additional insights regarding their role in tumor progression and potentially enabling novel avenues for immunotherapies.

## Results

### Single-cell profiling of T cells in CRC and adjacent normal colon

We analyzed 37,931 single T cells in CRC and adjacent normal colon from total 16 patients including its single-cell transcriptome, 23 cell surface marker expressions, and TCR sequences (Fig. 1a-c, and Extended Data Fig.1a). Viable TCRαβ^+^ CD3^+^ T cells were sorted and single-cell RNA sequencing (scRNA-seq) and TCR sequencing (TCR-seq) were performed^27^ (Fig. 1a and Extended Data Fig.1a). Cells from different normal colon/tumor samples or stimulated/non-stimulated fractions were resolved through labeling with distinct antibody hashtags^28^ (Fig. 1c,h, Extended Data Fig. 1C, Methods). After stringent quality filtering, we obtained 35,145 single T cell transcriptomes. TCR-seq analysis assigned annotated TCR sequences for 32,550 single T cells, of which 27,999 cells had paired TCR sequences (Methods).

**Fig. 1.**
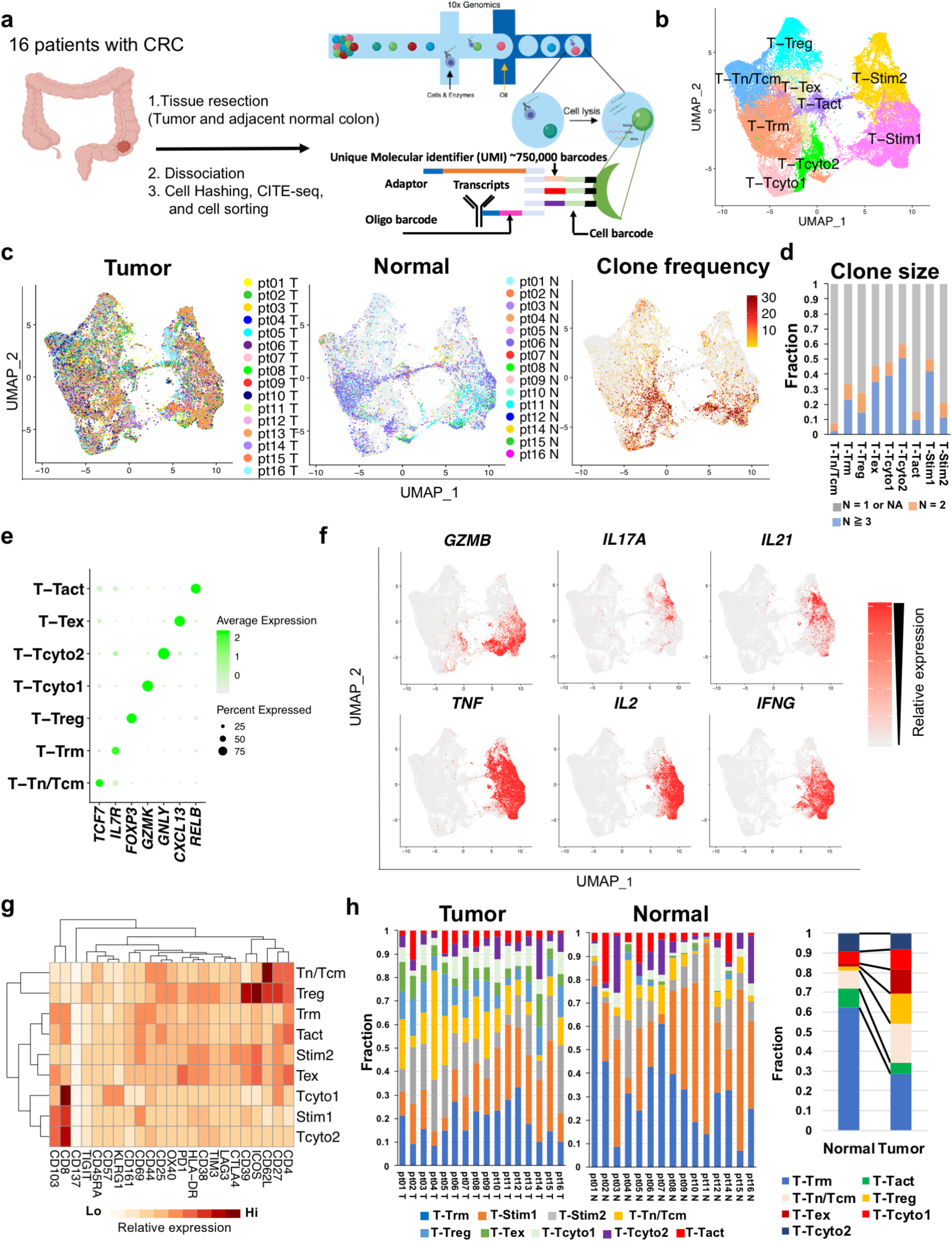
Characterization of T cells in CRC and normal adjacent colon by multiplexed scRNAseq. **a**, Workflow for preparation of multiplexed scRNA-seq and TCR-seq libraries using 10x Genomics platform. **b**, Uniform Manifold Approximation and Projection (UMAP) of 35,145 single T cells in CRC and adjacent normal colon from total 16 patients. Distinct clusters based on unsupervised clustering of single-cell transcriptomes are indicated. Unstimulated T cells populate within 7 clusters (left), stimulated cells T cells populate two clusters (right). The number of tumor-infiltrating T cells or normal cells within each cluster is indicated in Supplementary Table 4. **c**, Single T cells within tumor (left) or adjacent normal colon (middle) from each patient (as indicated) are plotted on the total UMAP in Fig. 1b. The number of tumor-infiltrating T cells or normal cells from each patient is indicated in Supplementary Table 4. Cells are colored based on TCR clone frequency (right), normalized to total number of T cells per patient sample. **d**, Bar graph depicting clonal composition of T cells within each cluster. **e**, Dot plot depicting relative expression of identifying marker genes for each non-stimulated cluster in Fig. 1b **f**, Heatmap of cytokine and effectors based on relative gene expression (as indicated) on the total UMAP in Fig. 1b **g**, Heatmap depicting average ADT signal in the CITE-seq antibody panel within each cluster. **h**, Bar graphs showing relative distribution of cells from each cluster (as indicated) within tumor (left) or normal adjacent colon (middle) of each patient (as indicated), and proportion of cells within each cluster from tumor or normal tissue (right).

Unsupervised clustering identified 9 distinct T cell clusters consisting of non-stimulated and stimulated clusters (Fig. 1b,e, Extended Data Fig. 1c). Stimulated T cell clusters (T-Stim1 and T-Stim2) that produced effector molecules and cytokines such as *GZMB, IL2, TNFA, IFNG, IL21* and *IL17A*, were composed primarily of CD8^+^ and CD4^+^ T cells, respectively (Fig. 1b,f, Extended Data Fig.1c,d).

The T_Tn/Tcm (naïve/central memory) cluster was enriched for markers associated with naïve T cell phenotypes (*TCF7, CCR7*, CD62L^+^, CD45RA^+^, CD27^+^). There was little clonal expansion present within this population (Fig. 1c,d). The T_Treg (regulatory T) cluster consisted primarily of CD4^+^ T cells, and was enriched for markers associated with Tregs (*FOXP3, BATF*, CD25^+^, CD62L^+^, CTLA4^+^, CD27^+^, ICOS^+^, TIGIT^+^, OX40^+^, TIM3^+^) (Fig. 1e,g, Extended Data Fig. 1b,d,e). Both T_Trm (resident memory T) and T_Tact (activated T) clusters were enriched for markers of Trm cells (CD69^+^, CD103^+^) (Fig. 1g, Extended Data 1e). The T_Trm cluster, the most diverse in terms of gene expression, consisted of both CD4^+^ and CD8^+^ T cells with distinct signature genes (*IL7R, JAML, ITGA1*), and was widely present in both normal colon and tumors (Fig. 1c,h, Extended Data Fig. 1b,d). The T_Tact cluster exhibited signatures of T cell activation (*RELB, NFKB2*, CD25^+^, CD69^+^, CD27^+^) (Fig. 1e,g, Extended Data Fig. 1e). The T_Tex (exhausted T) cluster was enriched for expression of genes (*CXCL13, HAVCR2*) and surface proteins (PD-1, ICOS) associated with T cell exhaustion or dysfunction^34,35^ (Fig. 1e,g, Extended Data Fig. 1b). Cells within this cluster were highly clonally expanded and co-expressed CD39 and CD103, consistent with tumor-infiltrating T cells with tumor-reactivity^36,37^ (Fig. 1g, Extended Data Fig. 1e).

We identified two clusters of cytotoxic T cells, T_Tcyto1 (cytotoxic T1) and T_Tcyto2 (cytotoxic T2), expressing signature genes of effector function and cytotoxicity (Fig. 1b,e, Extended Data Fig.1b). The T_Tcyto1 cluster was enriched for markers of effector memory T cells (Tem; *EOMES, GZMK*, KLRG1^hi^, CD57^hi^, CD27^hi^, CD44^+^), while the T_Tcyto2 cluster (*GNLY, GZMH, PRF1*) was enriched for expression of the resident memory marker CD103 (Fig. 1e,g, Extended Data Fig. 1b,d). Both clusters consisted primarily of CD8^+^ T cells with high clonal expansion, although CD4^+^ T cells were present in both populations (Extended Data Figs. 1d and 5a). Cells in the T_Tcyto1 cluster were relatively enriched within tumors compared to normal colon with high clonal expansion (Fig. 1c,d,h).

Normal T cells exhibited relatively less phenotypic diversity, and were essentially absent in T_Treg and T_Tex clusters (Fig. 1h, Extended Data Fig. 2d). In contrast, intratumoral T cells were widely distributed over all the clusters and each T cell type was observed in all the patients (Fig. 1c,h and Extended Data Fig. 2), suggesting that functional insights and gene signatures gleaned through this analysis are broadly applicable in human CRC (Fig. 1c,h).

**Fig.2.**
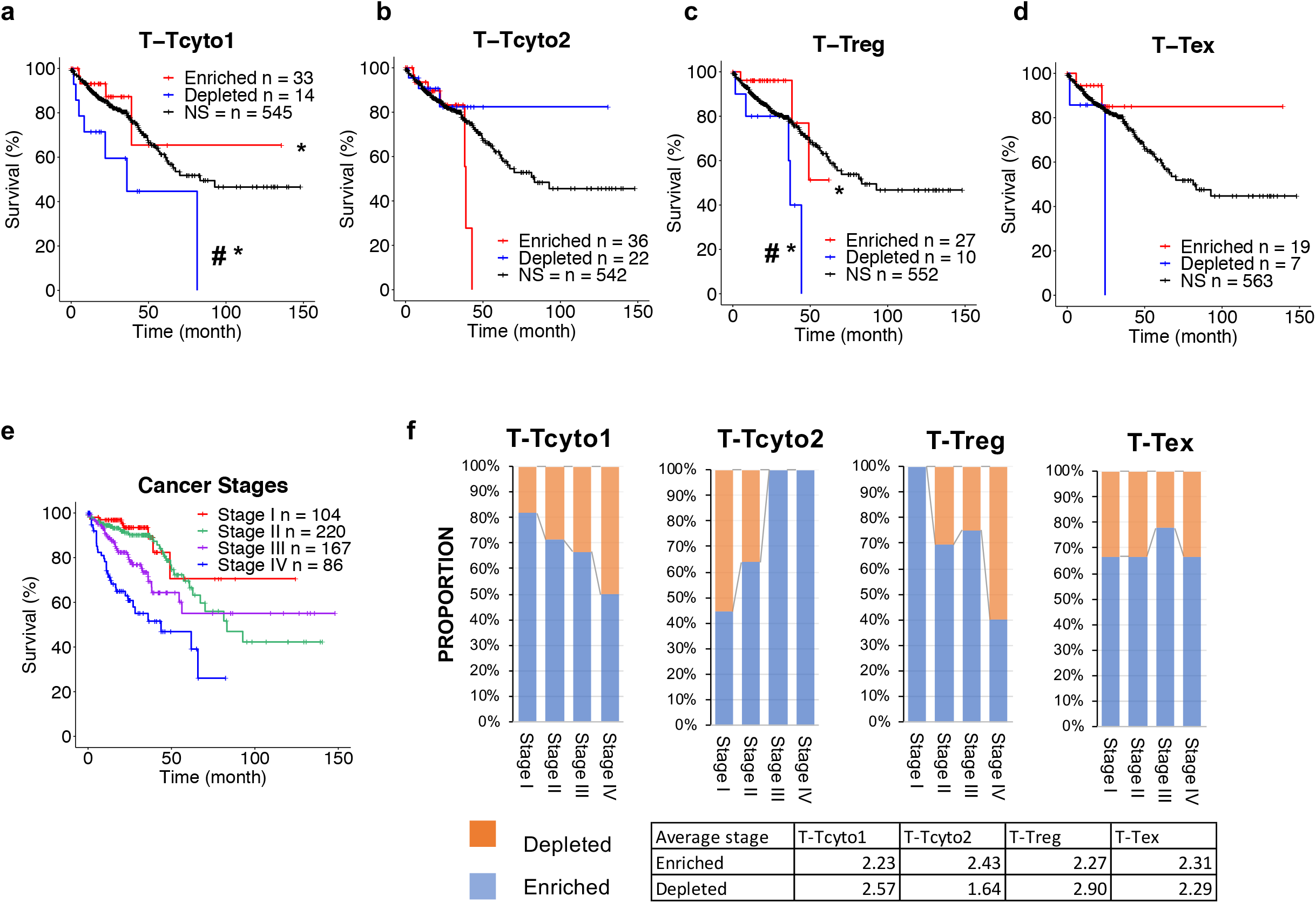
Clinical outcomes associated with T cell subtypes identified by scRNA-seq in CRC. (**a**-**d**) Kaplan–Meier curves of overall survival in the CRC TCGA cohort for patients enriched or depleted for the following gene sets using GSEA (Methods): T-Tcyto1 (**a**) T-Tcyto2 (**b**) T-Treg (**c**) or T-Tex clusters (**d**). ^#^denotes *P* < 0.05 (log-rank test) compared to non-enriched, non-depleted patients (curve at black). *denotes *P* < 0.05 (log-rank test) when enriched vs. depleted patients are compared. See detailed p-values (Supplementary Table 6). **e**, Kaplan-Meier curves of overall survival by clinical tumor-stage for the CRC TCGA cohort. **f**, Bar graph depicting relative proportion of patients by stage enriched or depleted for T-Tcyto1, T-Tcyto2 or T-Treg gene sets. Average stage of patients is indicated in table on bottom.

### Prognostic significance of distinct T cell populations with effector functions in CRC

To investigate the prognostic significance of distinct T cell subtypes, we applied semi-quantitative gene set enrichment analysis (GSEA) to The Cancer Genome Atlas (TCGA) database^31^ (Methods). We queried the top 30-40 differentially expressed genes in non-stimulated clusters with effector function (T_Tcyto1, T_Tcyto2, T_Tex, and T_Treg) within bulk RNAseq data from the CRC TCGA cohort (Supplementary Table 5). We subsequently compared survival between the statistically gene-enriched and -depleted patients through Kaplan-Meier analysis (Methods). CRC patients who were enriched with the T_Tcyto1 gene set (n= 33) showed a favorable prognosis compared to the depleted patients (n=14, *P* < 0.05) (Fig. 2a), whereas there was no statistically significant difference in survival between T_Tcyto2 gene-enriched (n=36) and depleted patients (n=22, *P* = 0.2) (Fig. 2b). These results suggest that T_Tcyto1 and T_Tcyto2 clusters contain functionally distinct T cells (mainly CD8^+^ T cells). In support of this, the proportion of the T_Tcyto1 gene-enriched patients decreased with late-stage cancer, while the proportion of T_Tcyto2 gene-enriched patients increased (*P* < 0.004; two-sided fisher exact test) (Fig. 2f). As expected, stage influenced survival (Fig. 2e). Thus, it is likely that the positive prognostic significance of the T_Tcyto1 cluster is at least partially dependent on its relative enrichment in early-stage cancer (Fig. 2f).

We also queried gene sets from the T_Tcyto1 and T_Tcyto2 clusters in gene expression data from the TCGA cohort of melanoma (Supplementary Table 5, Methods). In the melanoma cohort, positive clinical outcomes were associated with both gene sets from the T_Tcyto1 (*P* < 0.05) and the T_Tcyto2 cluster (*P* < 0.05) (Extended Data Fig. 3a,b). As expected, survival correlated with stage (Extended Data Fig. 3d). In contrast to CRC, the proportion of the gene-enriched patients did not correlate with tumor-stage in melanoma (Extended Data Fig. 3e). Moreover, there was higher overlap between gene-enriched patients from these clusters in melanoma (86 %; T_Tcyto1, 78 %; T_Tcyto2) compared to CRC, likely reflecting differences in cytotoxic T cell populations between these two cancers (Extended Data Fig. 3f,g).

**Fig. 3.**
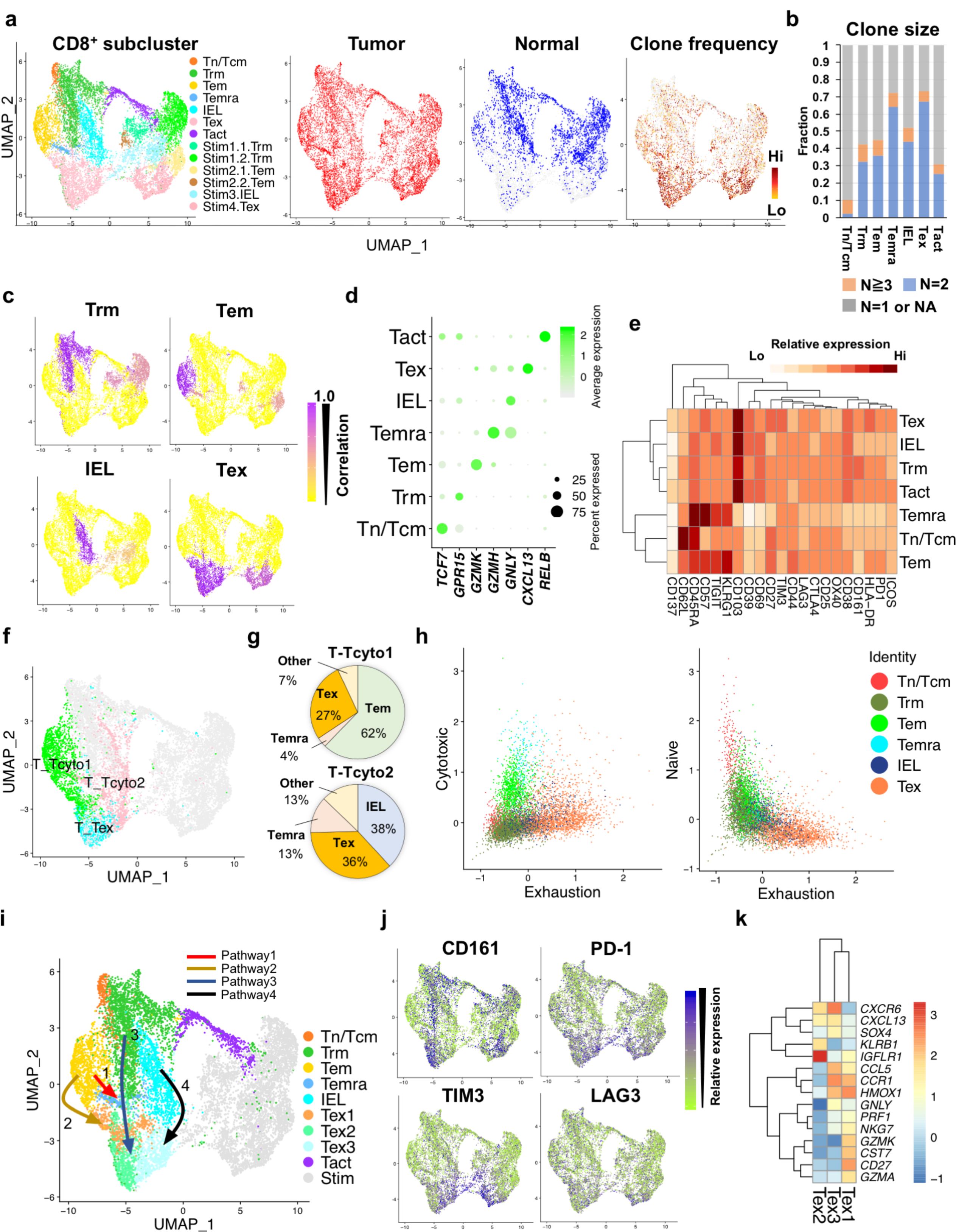
Dynamic phenotypic changes of CD8^+^ T-cell infiltrates in CRC. **a**, UMAP of 19,183 single CD8^+^ T cells in CRC and adjacent normal colon from 16 patients (left). Unstimulated T cells populate within clusters on left side, stimulated cells T cells populate clusters on the right side. Cells from tumor (red) or adjacent normal colon (blue) are indicated (middle). Cells are colored based on TCR clone frequency (right), normalized to total number of T cells per patient sample. **b**, Bar graph depicting clonal composition of T cells within each cluster. **c**, Pearson correlation map between each cluster in non-stimulated and stimulated cells, correlated based on 21 ADT signals. **d**, Dot plot depicting relative expression of identifying marker genes for each non-stimulated cluster in Fig. 3a. **e**, Heatmap depicting average ADT signal in the CITE-seq antibody panel within each non-stimulated cluster. **f**, Distribution of cells within the clusters T-Tcyto1, T-Tcyto2, T-Tex (in Fig. 1b) plotted on the CD8 UMAP. **g**, Pie graphs depicting composition of cells within the clusters T-Tcyto1 or T-Tcyto2 as a proportion of CD8 clusters CD8_Tem, CD8_Temra, CD8_IEL, CD8_Tex. **h**, Dot plots of cells from each CD8 non-stimulated cluster by naïve vs. exhaustion score or cytotoxic vs. exhaustion score (Methods). **i**, Distinct differentiation pathways of CD8^+^ T cells based on cell trajectory (relate to Extended Data Fig. 4f,g). **j**, Heatmap of ADT signals (as indicated) on the CD8 UMAP. **k**, Gene heatmap of differentially expressed genes within Tex subpopulations (Tex1-Tex3) in Fig. 3i.

The gene set from the T_Treg cluster also correlated with a positive prognosis (*P* < 0.05) in CRC, but had no significant correlation (*P* = 0.5) with prognosis in melanoma (Fig. 2c, Extended Data Fig. 3c). These findings are consistent with the preponderance of histologic studies that concluded that Tregs are associated with favorable outcomes in CRC^38,39^. Notably, relative enrichment of T_Treg genes in the patients was highest in early-stage cancer, indicating that the improved prognosis may be related to stage (Fig. 2f). Analysis of the prognostic value of the T_Tex gene set did not reach statistical significance (*P* = 0.1) in CRC (Fig. 2d).

### Distinct differentiation pathways of CD8^+^ T-cell infiltrates including prognostic effector memory and non-prognostic resident memory cells

To define T cells with improved granularity, we segregated CD8^+^ and CD4^+^ T cells in total 35,145 single cells based on ADT (antibody-derived tag) signal and transcriptome, and analyzed them separately (Extended Data Fig. 4a). Within 12,642 single CD8^+^ T cells, unsupervised clustering based on transcriptome identified 13 distinct clusters consisting of non-simulated and stimulated CD8^+^ T cells (Fig. 3a). Based on 21 ADT signals (except CD4 and CD8), we were able to link non-stimulated clusters to its cognate stimulated (Fig. 3c, Methods). Overall, all CD8 subtypes were observed within tumors, whereas normal cells were largely confined to the clusters CD8_Trm, CD8_IEL, and CD8_Tact (Fig. 3a).

**Fig. 4.**
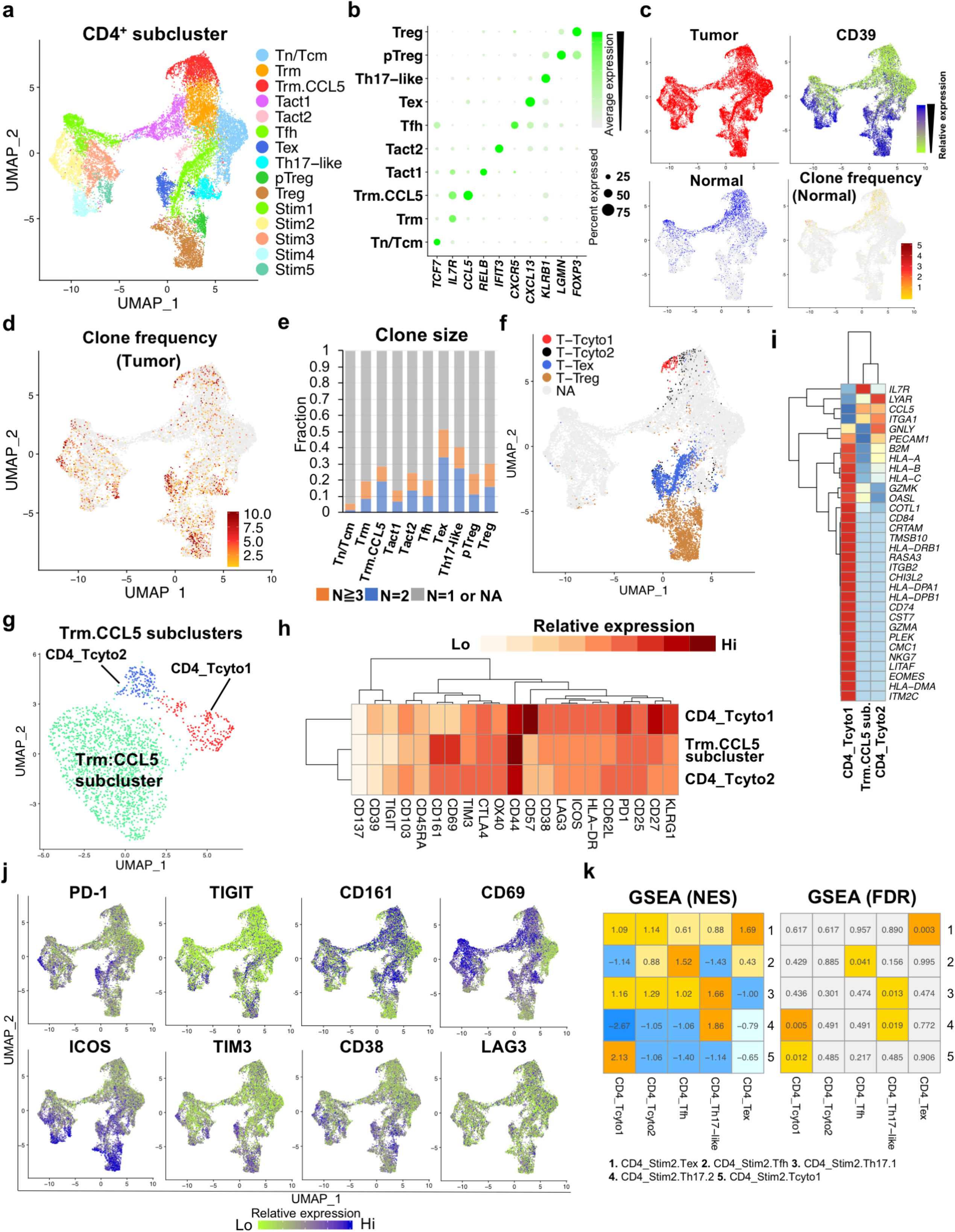
The functional and phenotypic diversity of single CD4^+^ T cells with prognostic significance in CRC. **a**, UMAP of 12,642 single CD4^+^ T cells in CRC and adjacent normal colon from total 16 patients. Unstimulated T cells populate within clusters on right side, stimulated cells T cells populate clusters on the left side. **b**, Dot plot depicting relative expression of identifying marker genes for each non-stimulated cluster in Fig. 4a. **c**, Cells from tumor or adjacent normal colon are indicated on the CD4 UMAP. Heatmap of CD39 ADT signal and heatmap of TCR clone frequency in normal cells. on the CD4 UMAP. **d**, Heatmap of TCR clone frequency in intratumoral cells on the CD4 UMAP. **e**, Bar graph depicting clonal composition of T cells within each cluster. **f**, Distribution of cells within the clusters of T-Tcyto1, T-Tcyto2, T-Tex, and Treg (identified in Fig. 1b) on the CD4 UMAP. **g**, Subclustering of CD4_Trm.CCL5 cluster. 3 subpopulations are labeled. **h**, Heatmap depicting average ADT signal in the CITE-seq antibody panel within each CD4_Trm.CCL5 subcluster shown in Fig. 4g. **i**, Heatmap showing relative gene expression of the genes indicated within each CD4_Trm.CCL5 subcluster shown in Fig. 4g. **j**, Heatmap of ADT signals (as indicated) on the CD4 UMAP. **k**, GSEA based on gene expression between non-stimulated clusters and stimulated subclusters (as indicated).

Within non-stimulated cells, the CD8_Tn/Tcm cluster contained cells with relatively higher *TCF7* expression and the least clonal expansion (Fig. 3a,b,d,e, Extended Data Fig. 4b,e). The CD8_Trm, CD8_IEL (intraepithelial lymphocyte), and CD8_Tact clusters all consisted predominantly of CD8^+^ Trm cells (CD69^+^, CD103^+^) (Fig. 3a,e). The CD8_Trm cluster (*GPR15, IL7R*) mainly contained cells with relatively higher expression of CD161, described to define a particular memory cell-type in CD8^+^ T cells including MAIT cells (CD161^hi^, *RORC, IKZF2*) expressing the semi-invariant TCR Vα7.2-Jα33/12/20^40^, which were present at low frequency (∼ 4 %) within this cluster (Fig. 3d,j, Extended Data Fig.4b, Supplementary Table 7). Cells within its cognate stimulated clusters (CD8_Stim1.1.Trm, CD8_Stim1.2.Trm) expressed cytokines including *TNF, IL2*, and *IFNG* (Extended Data Fig. 4d).

Cells within the CD8_IEL cluster expressed effectors (*GNLY, PRF1*) with expression of IEL (intraepithelial lymphocyte) gene signatures such as KIRs (Killer cell immunoglobulin-like receptors; *KIR3DL1/2*)^41^, and its cognate stimulated cluster (CD8_Stim3.IEL) contained cells expressing effector molecules, *GNLY, GZMB*, and *PRF1*. (Fig. 3d, Extended Data Fig. 4b,d). The CD8_IEL cluster overlapped with the T_Tcyto2 cluster, which accounted for 38% of cells within the cluster (Fig. 3f,g). The CD8_Tact cluster consisted of activated T cells (*NFKB2, RELB*) and contained both stimulated and non-stimulated cells (Fig. 3a,d, Extended Data Fig. 4b). The CD8_IEL and CD8_Tact clusters were associated with relatively high co-expression of CD39 and CD103, proposed to denote tumor-reactivity^36,37^ (Extended Data Fig. 4c).

The CD8_Tem cluster contained Tem cells (*EOMES, GZMK*, CXCR3, KLRG1^+^, CD57^+^, CD44^+^, CD27^+^) with relatively higher expression of naïve T cell markers (*CCR7, TCF7*), similar to “effector” T cells previously described in several types of tumors^9,42,43^ (Fig. 3d,e,h, Extended Data Fig. 4b, Supplementary Table 8). Cells within its cognate stimulated cluster, CD8_Stim.Tem, expressed *IL2, TNFA*, and *IFNG* (Extended Data Fig. 4d). Importantly, the CD8_Tem cluster largely overlapped with the positively prognostic T-Tcyto1 cluster, which constituted 62 % of the cells within this cluster (Fig. 3f,g).

The CD8_Temra cluster contained terminally differentiated Temra (effector memory CD45RA^+^) cells (*GZMH, GNLY*, CD45RA^hi^, KLRG1^hi^, CD57^hi^, TIGIT^hi^, CD27^−^) (Fig. 3d,e and Extended Data Fig. 4b). This cluster was enriched for “cytotoxic” signature genes (*FGFBP2, GZMH*) and contained highly clonally expanded cells^42^ (Fig. 3b,h, Extended Data Fig. 4b, Supplementary Table 8). The CD8_Tex cluster was enriched for Tex-cell markers (*HAVCR2*, CXCL13, PD-1^+^, TIM3^+^) and contained the most clonally expanded cells with co-expression of CD103 and CD39 (Fig. 3d,e,h, Extended Data Fig. 4b,c, Supplementary Table 8). Cells within its cognate stimulated CD8_Stim4.Tex cluster were capable of producing certain effectors (*PRF1, GZMB, IFNG)*, while relatively impaired for expression of *IL2* and *TNFA* (Extended Data Fig. 4d). Notably, the non-prognostic T_Tcyto2 cluster contained a large number of Tex cells (36%) as well as IEL(-like) cells (38%) (Fig. 3f,g).

To visualize the relationship between CD8^+^ T cell clusters, we used diffusion maps and ordered cells in pseudotime within non-stimulated clusters (Extended Data Fig. 4f, Methods). In combination with TCR sharing data, we found 4 differentiation pathways of CD8^+^ T cells within tumors, which culminated in terminally differentiated cell-types present within CD8_Temra or CD8_Tex clusters (Fig. 3i-k, Extended Data Fig. 4f,g). In one pathway (pathway 1), CD8_Tem cells differentiated into CD8_Temra cells. The other pathways culminated into three distinct Tex subpopulations: (pathway 2) CD8_Tem cells towards CD8_Tex1 subpopulation (*NKG7, GZMK, GZMA*), (pathway 3) CD8_Trm cells towards CD8_Tex2 subpopulation (CD161^hi^, *KLRB1, IGFLR1*), or (pathway 4) IELs towards CD8_Tex3 subpopulation (CD161^+^, *CXCR6, GNLY, CCL5*). Interestingly, cells within the positively prognostic T_Tcyto1 cluster were largely present in the CD8_Tem and Tex1 clusters with more effector function, while cells within the non-prognostic T_Tcyto2 cluster were present in the CD8_IEL and Tex3 clusters containing more dysfunctional cells (Fig. 3f,g,i). Thus, diverse types of CD8^+^ T cells (mainly prognostic Tem and non-prognostic Trm cells) transitioned to terminally differentiated cells through different pathways within tumors.

### Single CD4^+^ T-cell infiltrates with prognostic significance are phenotypically and functionally diverse

Unsupervised clustering of 19,183 single CD4^+^ T cells based on its transcriptome identified 10 distinct clusters consisting of non-stimulated and stimulated T cells (Fig. 4a,b, Extended Data Fig. 5a).

**Fig. 5.**
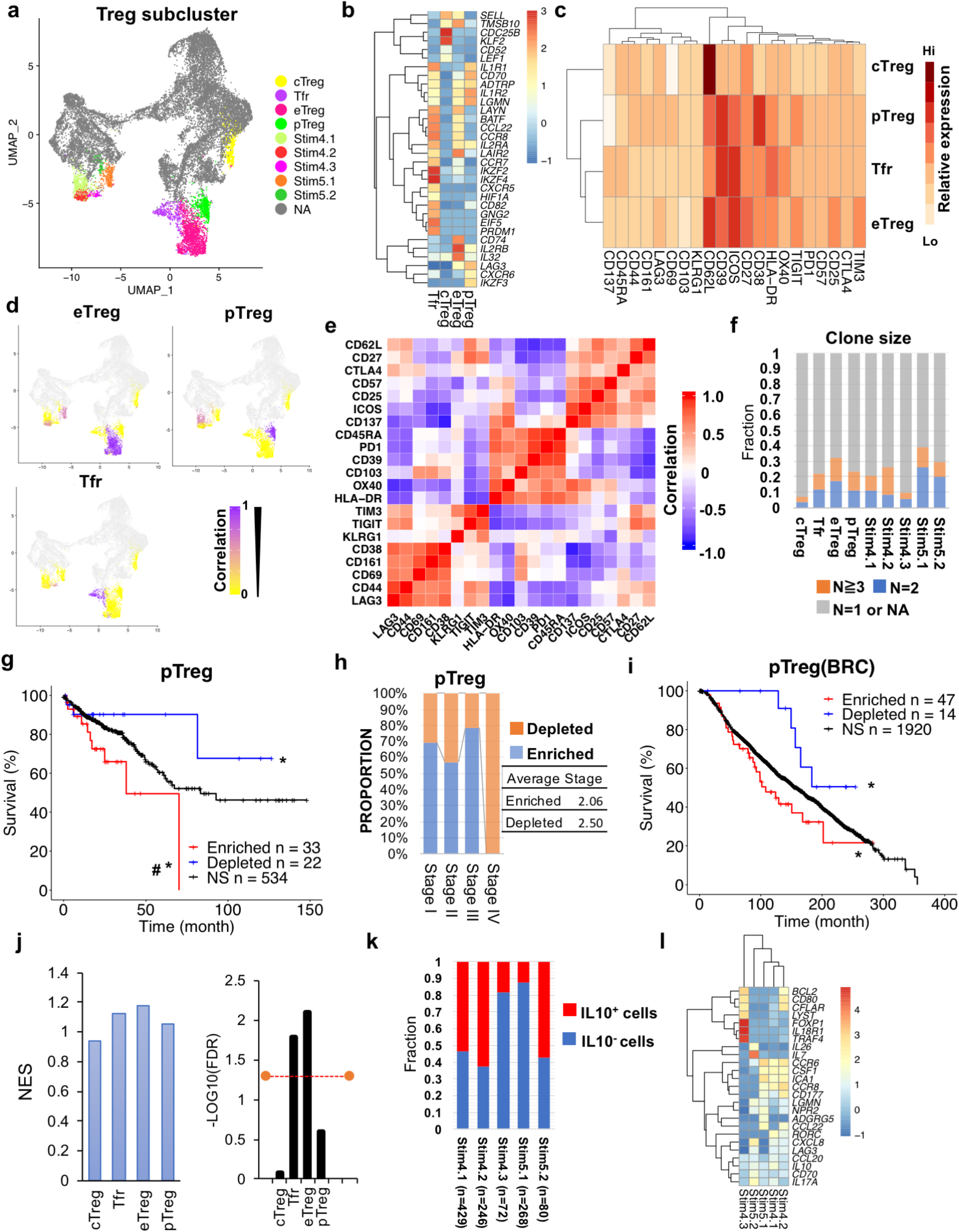
Prognostic significance of Treg subpopulations in CRC. **a**, Treg subpopulations on the CD4 UMAP from Fig. 4a. **b**, Heatmap of differentially expressed genes (as indicated) within Treg subpopulations. **c**, Heatmap depicting average ADT signal in the CITE-seq antibody panel within each non-stimulated Treg subpopulation. **d**, Heatmap of pearson correlation between non-stimulated and stimulated cells within each Treg subcluster, correlated based on 21 ADT signals. **e**, Heatmap showing pearson correlations between 21 ADT signals from each non-stimulated Treg subcluster. **f**, Bar graph depicting relative proportion of cells within each Treg subpopulation. (**g**,**i**) Kaplan–Meier curves of overall survival in the CRC (**g**) or BRC (**i**) TCGA cohort for patients enriched or depleted for the CD4_pTreg gene set by GSEA (Methods). ^#^denotes *P* < 0.05 (log-rank test) compared to non-enriched, non-depleted patients (curve at black). *denotes *P* < 0.05 (log-rank test) when enriched vs. depleted patients are compared. See detailed p-values (Supplementary Table 6). **h**, Bar graphs depicting relative proportion of patients by stage enriched or depleted for CD4_pTreg gene sets in CRC. Average stage is indicated on right. **j**, NES and FDR results from GSEA between the leading-edge genes (in Extended Data Fig. 7g) of the positively prognostic T-Treg cluster (in Fig. 2c) and signature genes in each Treg subpopulation. FDR threshold value indicated by the orange line is 0.05. **k**, Bargraphs depicting proportion of intratumoral *IL10*^+^ and *IL10*^−^ Tregs within each stimulated Treg subcluster. The number of cells within each cluster were indicated. **l**, Heatmap of differentially expressed genes in stimulated Treg subclusters.

Normal CD4^+^ T cells with a resident memory phenotype (CD69^+^, CD161^+^, *IL7R*) were present in non-stimulated clusters (CD4_Trm, CD4_Trm.CCL5, CD4_Tact1) and stimulated clusters (CD4_Stim1, CD4_Stim3), and were clonally expanded mostly within the CD4_Trm.CCL5 cluster with expression of CCL5, known to regulate proliferation and homeostatic maintenance of tissue Trm cells^44^ (Fig. 4a,c,d,j, Extended Data Fig. 5b). Stimulated normal cells expressed *IL2, IFNG, TNFA* and *IL10* and were associated with expression of Th1 marker (*Tbx21*; CD4_Stim1) or Tfh marker (*BCL6, IL21*; CD4_Stim3), whereas gene markers for Th17 and Treg cells, were mostly absent in normal cells (Extended Data Fig. 5c,d).

Intratumoral cells primarily contained effector CD4^+^ T cells expressing the activation marker CD39 (Fig. 4c,e). Many canonical genes that define effector CD4^+^ Th subsets were elicited through stimulation, including Th1, Th17, and Tfh signature genes (Extended Data Fig. 5c). The CD4_Stim2 cluster was further subdivided into 5 distinct subclusters by unsupervised clustering (Extended Data Fig. 5f). Through GSEA, we were able to link these subclusters with its cognate non-stimulated clusters and named them accordingly (Fig. 4k, Extended Data Fig. 5f,g).

Interestingly, CD4^+^ T cells within T-Tcyto1 and T-Tcyto2 clusters were observed in the CD4_Trm.CCL5 cluster, where further computational analysis identified 3 distinct subclusters including the CD4_Tcyto1 and CD4_Tcyto2 clusters predominantly containing T-Tcyto1 cells and T-Tcyto2 cells, respectively (Fig. 4f-I, Extended Data Fig. 5a). Cells within the CD4_Tcyto1 cluster (*NKG7, EOMES, GZMK, GZMA*, KLRG1^+^, CD57^+^, CD27^+^, PD-1^+^) were enriched within tumors and exhibited a similar phenotype to the CD8_Tem cluster (Fig. 4f-i, Extended Data Fig5a,e). Its cognate stimulated cluster (CD4_Stim2.Tcyto1) contained cells expressing effector cytokines, *IFN* and *IL2*, and chemokines (*CCL3, CCL4*) that can mediate T cell recruitment^45,46^ (Extended Data Fig. 5f,g). Likewise, cells within CD4_Tcyto2 exhibited a similar phenotype to the T_Tcyto2 cluster, and were not confined to tumors (Fig. 4f-i, Extended Data Fig. 5e).

The CD4_Th17-like cluster contained cells with Th17 cell signature genes (*KLRB1, RORA, CXCR6*) with high clonal expansion, and its cognate stimulated subclusters, (CD4_Stim2.Th17.1 and CD4_Stim2.Th17.2) contained cells expressing canonical Th17 genes (*RORC, IL17A*) (Fig. 4b, Extended Data Fig. 5b,c,f,g). Cells within the CD4_Stim2.Th17.1 cluster expressed genes associated with regulatory function (*IL10, ITGA2*)^47-49^, whereas cells within the CD4_Stim2.Th17.2 cluster expressed proinflammatory effectors (*IFNG, IL23R, PRF1*), described to be produced by pathogenic Th17 cells in autoimmunity^50^ (Extended Data Fig. 5g).

The positively prognostic FOXP3^+^ T cell population (T-Treg) was largely divided into 2 CD4 subpopulations (CD4_Treg, CD4_pTreg,) (Fig. 4a,b). The CD4_Treg cluster contained cells expressing distinct signature genes (*BATF, GATA3, IKZF2*) and high levels of ICOS (Fig. 5b,c, Extended Data Fig. 7a). The transcription factor Helios (encoded by the gene *IKZF2*) has been proposed to mark thymically-derived Tregs although currently controversial^51-53^. In contrast, Helios^−^ Tregs within the CD4_pTreg cluster exhibited a distinct phenotype (CD161^+^, CD69^+^, LAG3^+^, CD38^hi^) and shared genes (*IL26, KLRB1, IKZF3, LAG3*) with peripherally-derived CD4^+^ FOXP3^−^ T conventional (Tconv) cells, known as peripherally-derived Tregs (pTregs)^54-57^ (Figs. 4j and 5b,c,e, Extended Data Figs. 5h and 7a).

The CD4_Tfh cluster, containing PD-1^hi^ICOS^hi^CD27^hi^ *CXCL13*^+^ T cells with expression of naïve T cell markers (*CCR7, TCF7*), correlated with its stimulated clusters (CD4_Stim2.Tfh, CD4_Stim3) with expression of Tfh signature genes (*BCL6, CXCR5, IL21*) (Fig. 4b,h, Extended Data Figs. 5b,c,d, and 7). The CD4_Tex cluster (*CXCL13, GNLY, GZMA*, TIGIT^hi^, ICOS^hi^, CD27^−^) with expression of co-inhibitory molecules (PD-1^hi^, TIM3^hi^) and exhausted marker genes (*HAVCR2, TOX)*^40,58^ contained cells with high clonal expansion, and correlated with the stimulated CD4_Stim2.Tex cluster with expression of *IFNG* and *IL21* (Fig.4b-f,h, Extended Data Fig. 5b). The CD4_Tact2 cluster contained CD4^+^ T cells with high relative expression of type-1 interferon (IFN)-inducible genes (*IFIT3, MX1*, and *HERC5*)^59,60^ (Extended Data Fig. 5b).

We found that cells within the CD4_Tfh cluster with a more naïve phenotype (CD39^lo^, *CCR7, TCF7*) shared TCRs with effector cells within the several clusters (CD4_Tex, CD4_Th17-like, CD4_pTreg, CD4_Treg) (Fig. 4a,c,d,j, Extended Data Fig. 6). These results suggest that the CD4_Tfh cluster could contain a transitional population that gives rise to more differentiated effector CD4^+^ T cell populations. Through GSEA, we did not observe any correlation of CD4_Tfh, CD4_Th17-like, or CD4_Tex gene sets with clinical outcomes (Extended Data Fig. 6).

### CD38^+^ pTreg-infiltrates correlate with poor clinical outcomes in CRC

To investigate the relationship between non-stimulated and stimulated clusters, Treg populations were further subdivided by unsupervised clustering, resulting in 4 distinct non-stimulated and 5 distinct stimulated subpopulations. We designated non-stimulated subpopulations cTreg (central memory), pTreg (peripherally-derived), eTreg (effector), and Tfr (follicular) (Fig. 4a, Fig. 5a,b,c, Extended Data Fig. 7a). Based on 21 ADT signals, we were able to link these Treg subpopulations with its cognate subpopulations within stimulated subclusters (Stim4.1, Stim4.2, Stim4.3 within Stim4 cluster; Stim5.1, Stim5.2 within Stim5 cluster) (Fig. 5d).

Helios^+^ Tregs within the CD4_Treg cluster were subdivided into the CD4_eTreg and CD4_Tfr clusters (Fig. 5a, b). The CD4_eTreg cluster, containing cells highly expressing CD62L, ICOS, and CD25 with high clonal expansion (>30%), correlated with its cognate stimulated subclusters (CD4_Stim4.2 and CD4_Stim5.1) containing cells with expression of genes related to trafficking (*CCR6, CCL22*) (Fig. 5c,f,l). The CD4_Tfr cluster was phenotypically similar to the CD4_eTreg, but distinguished by a Tfr marker gene *CXCR5* (Extended Data Fig. 7a). Tfr cells also expressed the gene *PDLM1*, encoding the transcription factor Blimp-1, a marker of eTregs^61^, and had high relative expression of OX40 and HLA-DR (Fig. 5b,c). The CD4_Tfr cluster correlated with cognate stimulated CD4_Stim4.3 cluster with genes (*IL18R1, FOXP1, BCL2, TRAF4*) related to the activation and maintenance of Tregs^62-65^ (Fig. 5d,l).

The CD4_pTreg cluster was linked to cognate stimulated clusters CD4_Stim4.1 and CD4_Stim5.2, which expressed Th17 signature genes, *IL17A* and/or *RORC* and a higher level of *IL10* compared to other Treg clusters (Fig. 5b,k,l, Extended Data Fig. 7a). Through GSEA, the gene set from the CD4_pTreg was associated with poor clinical outcomes in not only the CRC but also the BRC cohort (Fig. 5g,i, Methods). Moreover, the gene set enrichment was observed in patients with early-stage cancer (except Stage IV), indicating that pTreg enrichment was independent of stage (Fig. 5g). Interestingly, there was substantial overlap between CD4_pTreg-enriched and CD4_Th17-like-enriched patients within the CRC cohort (76%; 25 of 33 enriched patients), but no overlap between CD4_pTreg-enriched and CD4_Th17-like-depleted patients (Extended Data Fig. 7c). In addition, pTregs and Th17-like clusters shared T cell clones, although TCR sharing was not confined to these clusters (Extended Data Fig. 7f). These results indicate that negatively prognostic pTregs and Th17 cells in CRC are functionally and phenotypically linked, and enriched in the same patients.

Although neither CD4_eTreg nor CD4_Tfr gene sets were associated with a favorable prognosis CRC, leading-edge genes within the positively prognostic T_Treg cluster were relatively enriched in eTreg and Tfr cells, but not pTregs (Fig. 5j, Extended Data Fig. 7d,e,g). Also, both eTreg and Tfr cells had significantly higher frequency of TCR sharing, compared to pTregs (*P* < 0.0001) (Extended Data Fig. 7b). These results suggest that Helios^+^ Tregs, as opposed to Helios^−^ Tregs, account for Tregs associated with favorable outcomes in CRC. Taken together, these data demonstrate substantial phenotypic and functional heterogeneity of Tregs within CRC, and provides clues on how these cells influence outcomes.

## DISCUSSION

Prior studies, performed through histology or analysis of bulk RNA, have demonstrated a relationship between T-cell infiltrates and clinical outcomes in CRC^1,2,8,11,66^. However, as a relatively limited number of genes were assessed in these studies, the resolution to assess the significance of distinct T cell subtypes was limited.

Our single-cell analysis delineated the functional and phenotypic diversity of TILs and its prognostic significance in CRC. For CD8^+^ T-infiltrates in CRC, we found that KLRG1^+^ *GZMK*^+^ Tem cell population could differentiate into 2 different cell-types. One contained a Tex population at a less dysfunctional transcriptional state (CD39^+^, CD103^+^, *GZMA, GZMK, PRF1*), enriched in early-stage cancer. The other contained CD45RA^+^ Temra cells with senescent cell markers (KLRG1^hi^, CD57^hi^)^67^. Moreover, 2 different cell-types of non-prognostic Trm cells (IELs and CD161^+^ T cells) were relatively enriched in late-stage cancer. These cells also differentiated into 2 different Tex populations primarily within tumors. Of these four pathways, the differentiation pathway including *GZMK*^+^ Tem and the more functional Tex subcluster accounted for a majority of positively prognostic cytotoxic T cells. These findings imply that factors driving distinct pathways of differentiation within CRC may be a key for anti-tumor immunity and a therapeutic avenue. Further, our data indicate that CD8^+^ T cell populations within CRC evolve with stage, transitioning from effector to IEL phenotypes with tumor progression.

Compared to CD8^+^ T cells, CD4^+^ T cell subtypes within tumors exhibited more complexity and diversity. Notably, we found that the prognostic Tem cells contain cytotoxic CD4^+^ T cells (but not Th1 cells) with a similar phenotype of CD8^+^ T cells. As a recent report indicated anti-tumor reactivity of cytotoxic CD4^+^ T cells in bladder cancer, further investigation could detail their role in CRC.

Although T-bet^+^ T cell infiltration was reported to be associated with a favorable prognosis in CRC^1,2^, our single-cell analysis showed substantial diversity in CRC, with expression of T-bet observed in not only Th1 cells but also Th17 cells and Tregs^68,69^. T-bet^+^Th17 cells were primarily enriched within tumor rather than canonical Th1 cells. We were not able to stratify such gene sets-enriched or -depleted patients from bulk-seq data of TCGA cohort (possibly due to loss of effector gene expression without any stimulation); it is worthwhile validating the contribution of these specific effector T cells to clinical outcome for future.

*CXCL13*^+^CD4^+^ T cells including Tfh and CD4^+^ Tex cells were mainly observed within tumors and both populations highly expressed PD-1. In addition, cells within the CD4_Tex cluster were enriched in MSI-H patients (53 %; 16 of 30 enriched patients) consistent with the prior study suggesting these cells could respond to PD-1 blockade^9^. Based on GSEA analysis, CD4_Tex-cell infiltrates were not predictive of prognosis.

For tumor-infiltrating Tregs in CRC, our data showed that almost all Tregs were specifically enriched within tumors (but not normal colon) and highly expressed the activated markers of CD39 and ICOS. Phenotypically, these Tregs were further subdivided into Helios^+^ Tregs and Helios^−^ Tregs (pTregs). pTreg signature genes were associated with a poor prognosis, while the gene set from the T-Treg cluster (before subclustering) was associated with a favorable prognosis. These results could be explained by our data that gene signatures of Helios^+^ Tregs accounted for a large number of prognostic leading-edge genes of the T-Treg cluster, suggesting that Helios^+^ Tregs rather than pTregs could correlate with positive clinical outcomes.

pTregs exhibited a Th17-cell phenotype with expression of *IL17A, RORC*, and *IKZF3* and shared TCRs with cells within the CD4_Th17-like cluster, although Th17(-like) cell-infiltrates were not associated with clinical outcomes. These results suggest that IL-17A^+^ pTregs rather than Th17 cells itself, positively involve tumor progression, and therefore may represent a target of immunotherapy. In support of this, no studies to date have considered the cellular source of IL-17, although IL-17 has been correlated with poor outcomes in CRC^9,11^. pTregs also showed a unique phenotype and function with higher expression of CD38 and LAG3 compared to Helios^+^ Tregs, and potentially highly produce IL-10. It is intriguing that Daratumumab, a monoclonal antibody used to treat myeloma, has been shown to target a CD38^+^ Treg subpopulation^70^.

## Acknowledgement

We thank patient volunteers and providers in the Departments of Surgery, Medicine, Pathology and the Herbert Irving Comprehensive Cancer Center at Columbia University Irving Medical Center for assistance in obtaining patient samples. We thank Hanina Hibshoosh and tissue bank for processing of patient samples. Erin Bush and Izabella Krupska within the JP Sulzberger Columbia Genome Center helped with single-cell analysis. We thank Michael Kissner and the flow cytometry cores within the Columbia Stem Cell Initiative and the Columbia Center for Translational Medicine (S10OD020056). This work was supported by the Damon Runyon Cancer Research Foundation, The Louis V. Gerstner Jr. Scholars Program, and K08 DK100738 (Han). We thank members of the Han lab and Yan lab for comments and suggestions.

## Author contributions

Study Concept and Design, K.M., A.H.; Patient and Sample Acquisition, K.M., S.L., M.I., V.R., A.M.A., S.A.L., P.R.K., K.S.Y., P.E.O, and A.H.; K.M., A.H. analyzed Data; Performed and designed experiments, K.M., S.L., K.S., M.S., P.S; Computational Analysis, K.M., A.K., P.H., A.M.B. and P.A.S; Wrote manuscript, K.M., N.S. A.H.; Study oversight, A.H.

## Supplementary Data

Extended Data Figures and Supplementary Tables are available at Supplementary Materials.

## Methods

### Human colon biopsy and single-cell dissociation

Tumors and adjacent normal colon tissues from the patients, diagnosed with colorectal adenocarcinoma at Columbia University Irving Medical Center, were collected with informed consent, as approved by the Columbia University Institutional Review Board (IRB). All protocols were performed in accordance with the guidelines provided by the IRB. All patients were treatment naïve at time of sample acquisition (Supplementary Table 1).

Fresh tissue was collected the same day of surgery. Tissue was dissociated into single-cell suspensions with enzymic digestions using a gentleMACS dissociate per protocol (Miltenyi Biotec). Single-cell suspensions were cryopreserved (10% DMSO) and stored in a liquid nitrogen.

### Antibody-oligo conjugates

Antibodies for CITE-seq and Cell Hashing were purchased from BioLegend, and were covalently conjugated to barcoded oligos by click-chemistry tools in accordance with the protocol of CITE-seq Hyper Antibody-Oligo Conjugation (https://cite-seq.com/protocol/). The list of antibodies, clones and barcodes are available in Supplementary Table 2.

### T cell isolation, T cell stimulation and cell sorting from CRC and adjacent normal tissues

Cryopreserved single-cell suspensions were thawed at 37 degrees, and single T cells were isolated by percoll (Millipore-Sigma) density gradient (20/40/80%) centrifugation (2200 rpm for 30 min). T cells were rested in complete RPMI in 96 well plates overnight. Next day, T cells from each patient were split into two wells, incubated with or without PMA/Ionomycin for 3h. After 3 washes, cells were incubated for 10 min with Fc receptor block (BioLegend) to block nonspecific antibody binding, each sample was stained with Cell hashing antibody cocktail for 30 min. After 3 washes, cells were stained with CITE-seq antibodies and FACS antibody cocktails for 30 min (Supplementary Table 2). Viable T cells were sorted by FACS Aria II by gating DAPI^−^ ZombiAqua^−^ CD3^+^ TCRαβ^+^ T cells (Extended Data Fig. 1a).

### Single cell 5’ transcript, TCRαβ enriched, and Cell hashing/CITE-seq library preparation on the 10x chromium, and sequencing

The scRNA-seq, scTCR-seq and HTO/ADT-seq libraries as mentioned below were prepared using the 10xChromium single cell 5’ library and gel bead Kit, per manufacturer’s instructions. In brief, oligo-conjugated antibody-tagged single cell suspensions were loaded into the Chromium^™^ controller to make nanoliter-scale droplets with uniquely barcoded 5’ gel beads called GEMs (gel bead-in emulsions). After GEM-RT, GEMs were cleaned up by Dyna beads MyOne^™^ Silane beads. At the cDNA amplification step, HTO/ADT additive oligos were added to amplify oligos derived from Cell hashing and CITE-seq antibodies. The products were size separated with SPRI Beads (Beckman Coulter): < 300 nt fragments containing the oligos derived from CITE-seq antibodies and Cell Hashing, and > 300 nt fragments containing cDNAs derived from cellular mRNA. HTO/ADTs were amplified using specific primers that append P5 and P7 sequences for illumina sequencing.

The cDNA was then pooled for downstream processing and library preparation according to the manufacturer’s instructions. The 5’ transcript library was sequenced with Illumina Novaseq. The HTO/ADT libraries were sequenced with Illumina Miseq or Nextseq. The single cell TCR enriched library was sequenced with Illumina Miseq using 150 paired-end reads.

### Data processing of HTO/ADT libraries

We counted antibody-derived-tags (ADTs) or Cell Hashing tags (HTOs) in raw sequencing reads and built a count matrix using CITE-seq-Count as previously described, available at (https://github.com/Hoohm/CITE-seq-Count). For HTO quantification, each HTO count was normalized by total counts of HTOs per cell barcode (total UMI counts were set at 100). Then, cell barcode identity was assigned to the most highly expressed HTO, otherwise assigned to “negative” or “doublet”, available at (https://github.com/akornberg/data_analysis_ak). Each CITE-seq UMI count was normalized by the average of total UMI counts from all cell barcodes, and then a count matrix was set into the Seurat as the ADT data, which was normalized based on the Seurat package, CLR normalization, and scaled.

### Data processing of scRNA-seq libraries

The total reads from scRNA-seq were aligned to GRCh38 reference genome and quantified using cellranger count (10x Genomics, version 3.1.0). In brief, only cells which passed the threshold set by the pipeline (more than 500 UMIs), were considered for further analysis. On average, we obtained reads from 1,862 genes per cell (median: 1,572) and 4,908 unique transcripts per cell (median: 4,178). The analysis of the created filtered feature-barcode matrices was performed using Seurat (version 3.1.3), available at (https://satijalab.org/seurat/).

### Data processing of single-cell TCR-seq libraries

The total reads from scRNA-seq were aligned to GRCh38 reference genome and consensus TCR annotation was conducted by cellranger vdj (10x Genomics, version 2.1.0). In total, 86 % (27,999 out of 32,550 cells) of annotated T cells were assigned a TCR (TRB and/or TRA), and 20,875 clonotypes were detected.

### Dimension reduction and clustering

For the downstream analysis, four multiple scRNAseq datasets (seurat objects) with different individuals and different conditions (Supplementary Table 3) were created by the Seurat, in which the metadata for HTO assignment (patients, stimulated or non-stimulated cells), normalized ADT counts (23 CITE-seq antibodies), and TCR clonotypes and sequences were combined into the datasets using the function AddMetadata and CreateSeuratobject, and were integrated using an anchor-based single-cell data integration method in Seurat^71^. In the method, the batch effects among the samples in four different datasets were normalized using the function the SCtransform in Seurat. Greater than 10% mitochondrial RNA content were excluded. Variable feature genes were set at 5,000 genes. Once the datasets were integrated, the data was used input into a principal component analysis (PCA) on the basis of variable genes. The same principal components were used to generate the UMAP projections. Clusters were identified using shared nearest neighbor (SNN)-based clustering. For the clustering analysis, the function RunUMAP, FindClusters, and FindNeighbors in Seurat were used, in which “dims” or “resolution” were set at between 10 and 30 or between 0.1 and 0.5, respectively.

### Analysis of TCR sharing and clone frequency

Identical TCR clonotypes (regarded as clonal identifier) among single-cells were counted through the Seurat metadata. Cells missing an identifier were considered as not observed (shown as NA). TCR sharing between two distinct clusters or different cell states on the UMAP, was assessed by the counts of identical TCRβ clonotypes. After CD4^+^/CD8^+^ single-cell separation, clone frequency in each single-cell was calculated as a relative value of TCRβ clonotype counts to total single-cells from each patient (TILs were separated from normal cells). Code is available at (https://github.com/akornberg/data_analysis_ak).

### Gene-gene Correlation Analysis and gene signature scoring

The stem-like, exhausted and cytotoxic signature genes were selected from variable genes across all CD8 T cells, which were computed using the Seurat: Those signature genes consisted of the top 20-30 genes with the highest correlation with the reference genes *TCF7, HAVCR2*, or *FGFBP2*, respectively as previously described^42^ (Supplementary Table 8). Individual cells on the CD8 UMAP (non-stimulated) were scored by the average of the selected gene expression, and shown as dot plots.

### Gene set enrichment analysis (GSEA) using TCGA cohort

Gene expression data of the TCGA cohort were obtained from cBioPortal (https://www.cbioportal.org/). The TCGA cohort data of CRC, BRC, and melanoma were downloaded from Colorectal Adenocarcinoma (TCGA, PanCancer Atlas), Breast Cancer (METABRIC), and Skin Cutaneous Melanoma^72,73^ (TCGA, PanCancer Atlas). Gene expression was normalized by log2-transformation and an average z-score across patient samples was calculated per gene. For each patient, genes were ranked by the average z-score in each gene. Regarding original gene sets in distinct T cell subsets delineated by unsupervised clustering, genes were ranked by differential expression test which was carried out using Seurat and the top genes (∼ 100 genes) were selected (min.diff.pct was set at > 0.25) after removing specific non-coding RNA such as lincRNA and miRNA, and genes linked with poorly supported transcriptional models (annotated with the prefix “AP-“) (Supplementary Table 5). The correlation between the gene sets and bulk databases of the TCGA cohort was calculated using pre-ranked gene set enrichment analysis method fgsea package (https://github.com/ctlab/fgsea).

### Prognosis prediction

The CRC, melanoma, or BRC TCGA cohort were grouped on the basis of GSEA, respectively. One group (NES, normalized enrichment score, >0 & a BH-adjusted p-value < 0.05) was categorized into “Enriched”. The other (NES <0 & a BH-adjusted p-value < 0.05) was assigned into “Depleted”. Kaplan-Meier survival curves were generated and log-rank test for two groups was performed using the R package survival (3.1.8).

### Diffusion map trajectory analysis

The Seurat object including the cluster CD8_Tn/Tcm, CD8_Trm, CD8_Tem, CD8_Temra, CD8_IEL, and CD8_Tex, which were characterized by unsupervised clustering in Seurat (Fig. 3a), was subjected into single-cell trajectory analysis using the Monocle (2.14.0), available at (http://cole-trapnell-lab.github.io/monocle-release/). RNA expression data from the Seurat object was converted into Monocle as a sparse matrix. The meta data from the Seurat object (total single cells in each patient, and cluster annotation) were incorporated as “phenodata” in Monocle. Then, newCellDataSet was created with a function “expressionFamily = negbinomial.size()”. In order to infer a single-cell trajectory, the total 650 genes (Supplementary Table 9) which were assembled from each cluster in the Seurat object using differential expression test, were chosen and then the cells were ordered after dimension reduction with “DDR-tree” function, and visualized.

### Gene or ADT expression heatmap and correlation analysis

For single-cell gene heatmaps, the representative marker genes were selected from differentially expressed genes within each cluster that was identified by unsupervised clustering. Gene heatmaps between cells within each cluster was created using the function DoHeatmap in Seurat (Extended Data Figs. 4b and 5b).

For ADT heatmaps and gene heatmaps among subclusters, each average ADT signal of cells within each cluster or average expression of each differentially expressed gene in each subcluster was calculated using the function AverageExpression in Seurat. Then, heatmap of each average ADT signal or gene expression was shown in selected clusters using the R package pheatmap (1.0.12).

The relationship between stimulated and non-stimulated clusters, was assessed by a statistical correlation between gene expression or average ADT signals in single-cells. In brief, for ADT correlation analysis, a matrix including each average ADT signal in stimulated and non-stimulated clusters was created and its correlation was examined by Pearson correlations. For gene expression correlation analysis, the representative marker genes were selected from differentially expressed genes within each non-stimulated cluster, respectively. And a matrix including each average variable gene expression in each stimulated cluster was created, respectively. Then, a correlation between marker genes in non-stimulated clusters and average gene expression in stimulated clusters was examined by GSEA using the R package fgsea.

## Code and Data Availability

The accession numbers for the sequencing results reported in this paper are **ArrayExpress**: E-MTAB-9455 (under revision). Fastq files from each sequencing lane (L1, L2) and corresponding to each fragment end (R1, R2) were concatenated into a single file to generate 2 fastq files (R1, R2) per biological replicate. Code is available at the links as indicated (Methods).

