## Extended Data Files for "The functional and phenotypic diversity of single T-cell infiltrates in human colorectal cancer as correlated with clinical outcome"

**Extended Data Fig.1 Landscape of single T cells in CRC and adjacent normal colon characterized by its transcriptome and expression of 23 cell surface proteins.**

**a**, Gating strategy for viable $CD3^{+}TCR\alpha\beta^{+}$ T cell sorting from dissociated cells. Single lymphocytes were gated based on scatter. Viable DAPI negative, Aqua LIVE/DEAD negative, $CD3^{+}TCR\alpha\beta^{+}$ cells are sorted and single-cell sequencing was performed. Sorting strategy was validated through staining with fluorescently-labeled CD4/CD8 antibodies that do not compete with ADT clones. Data are representative of four independent experiments.

**b**, Heatmap of differentially expressed genes between non-stimulated cells within each cluster (as indicated).

**c**, Distribution map of non-stimulated and stimulated cells on the total UMAP in Fig.1b. Unassigned cells are indicated in yellow.

**d**, Heatmap of CD4 or CD8 ADT signal on the total UMAP. Scale indicated on right.

**e**, Each ADT signal in the CITE-seq antibody panel on the total UMAP. Arrows indicate cell populations expressing rare markers.

**Extended Data Fig. 2 Cell composition of each patient within tumors.**

**(a-c)** Distribution map of intratumoral cells in each patient by tumor-stage.

**d,** Proportion of intratumoral or normal cells within each cluster (as indicated) and its comparison. ***, *P*-value (Wilcoxson-test) $<0.01$; * $P <0.05$; n.s. (not significant)

**Extended Data Fig. 3 Clinical outcomes associated with T cell subtypes identified by scRNA-seq in melanoma.**

**a-c**, Kaplan–Meier curves of overall survival in the melanoma TCGA cohort for patients enriched or depleted for the following gene sets using GSEA (STAR Methods): T-Tcyto1 (**a**), T-Tcyto2 (**b**), or T-Treg clusters (**c**). #denotes $P < 0.05$ (log-rank test) compared to non-enriched, non-depleted patients (curve at black). *denotes $P < 0.05$ (log-rank test) when enriched vs. depleted patients are compared. See detailed p-values (Supplementary Table 6).

**d**, Kaplan-Meier curves of overall survival separated by clinical tumor-stage for the melanoma TCGA cohort.

**e**, Bar graphs depicting relative proportion of patients by stage enriched or depleted for T-Tcyto1 (**a**), T-Tcyto2 (**b**). Average stage of patients is indicated in table on bottom.

**f,g** Venn diagram showing the number of patients enriched for T-Tcyto1, T-Tcyto2, or both (center at light blue) in melanoma (**f**) and CRC (**g**).

**Extended Data Fig.4 Functional properties and relationships between distinct CD8$^+$ T cell subtypes within tumors.**

**a**, Workflow of CD8$^+$/CD4$^+$ T cell separation based on ADT signals and transcriptome from total 35,145 single cells.

**b**, Heatmap of differentially expressed genes (as indicated) between cells within each CD8 non-stimulated cluster.

**c**, Heatmap of CD103 or CD39 ADT signal on the CD8 UMAP.

**d**, Heatmap of cytokine and effector genes (as indicated) on the CD8 UMAP.

**e**, Heatmap of the *TCF7* gene on the CD8 UMAP (top). *TCF7* mRNA expression levels are shown in density plot (bottom left). Bar blot depicts number of TCR clone cells with expression of high or low level of *TCF7* (separated by mean-value in density plot).

**f**, Pseudotime diffusion maps depicting differentiation of non-stimulated CD8$^+$ T cells. Cells colored by clusters (upper left) and pseudotime (lower left). Plots on right show normal or intratumoral cells from CD8 non-stimulated clusters, respectively.

**g**, TCR sharing between distinct cell states in Extended Data Fig. 4f. TCR clone cells were investigated between distinct clusters as indicated.

### Extended Data Fig. 5 Characterization of effector CD4$^+$ T cells within tumors.

**a**, Distribution map of non-stimulated CD4$^+$ T cells (in Fig. 4a) on the total T cell UMAP (in Fig. 1b). Arrows indicate CD4$^+$ T cells within the T-Tcyto1 and T-Tcyto2 clusters.

**b**, Heatmap of differentially expressed genes between cells in each CD4 non-stimulated cluster.

**c**, Heatmap of marker genes for Th1, Th17, or Tfh cells on the CD4 UMAP.

**d**, Heatmap of cytokine and effector genes (as indicated) on the CD4 UMAP.

**e,** Proportion of normal cells or intratumoral cells within each subcluster of CD4_Trm.CCL5 cluster for each patient (n=16) and its comparison. *** *P* (Wilcoxon-test) <0.01; * *P* <0.05; n.s.(not significant)

**f**, Distribution of cells within each CD4_Stim2 subcluster on the CD4 UMAP.

**g**, Heatmap of differentially expressed genes (as indicated) for cells within each CD4_Stim2 subcluster.

**h,** Comparison of ADT signal (as indicated) between CD4_pTreg cluster and CD4_Treg cluster cells. *** *P* (Wilcoxon-test) <0.0001

### Extended Data Fig. 6 Lineage and prognostic significance of peripheral CD4$^+$ T cells in CRC.

**a**, Heatmap of Tfh T cell clonotypes among the CD4 non-stimulated and stimulated clusters.

**b**, All expanded T cell clonotypes from cluster CD4_Tfh on the CD4 UMAP.

**c**, Kaplan–Meier curves of overall survival in the CRC TCGA cohort for patients enriched or depleted for CD4_Tfh, CD4_Tex, or CD4_Th17-like gene sets by GSEA (Methods).

**d**, Bar graphs depicting relative proportion of patients by stage enriched (blue) or depleted (orange) for the gene sets. Average stage is indicated on right.

**e**, Comparison of CD39 expression in between CD4_Tfh and other effector clusters as indicated. *P*-values (Wilcoxon-test) were indicated.

### Extended Data Fig. 7 Characterization of Treg subpopulations in CRC.

**a**, Relative gene expression of marker genes (as indicated) for pTreg, eTreg, or Tfr cells on the CD4 UMAP (in Fig. 4a).

**b**, eTreg TCR clonotype frequencies shared with Tfr or pTreg; *** $P < 0.0001$ (Wilcoxon-test).

**c**, Venn diagram showing the number of patients enriched for Th17 and pTreg genes (top). The same is shown for the number of patients depleted for Th17 and enriched for pTreg (bottom).

**d**, Kaplan-Meier curves of overall survival in the CRC TCGA cohort for patients enriched or depleted for CD4_Tfr or CD4_eTreg gene sets by GSEA (Methods).

**e**, Bar graph depicting relative proportion of patients by stage enriched (blue) or depleted (orange) for Tfr or eTreg subclusters. Average stage is indicated on right.

**f**, All expanded T cell clonotypes present in Tfr (left), eTreg (right) or pTreg (bottom) subsets on the CD4 UMAP.

**g**, Leading-edge genes of the patients enriched for the positively prognostic T-Treg cluster in Fig. 2c.

**Supplementary Tables**

Table 1. Patient characteristics

Table 2. Antibody clones and barcodes for CITE-seq and cell hashing

Table 3. Experimental datasets for scRNA-seq, TCR-seq, and CITE-seq

Table 4. Cluster cell number by patient and sample

Table 5. Gene sets for GSEA (related to Fig. 2 and Extended Data Fig. 3)

Table 6. P-values for Kaplan Meier analysis

Table 7. MAIT cell distribution in the CD8 UMAP

Table 8. Gene lists highly correlated with marker genes *TCF7*, *FGFBP2*, or *HAVCR2* (related to Fig. 3h)

Table 9. Gene lists for cell trajectory analysis (related to Fig. 3i and Extended Data Fig. 4f)

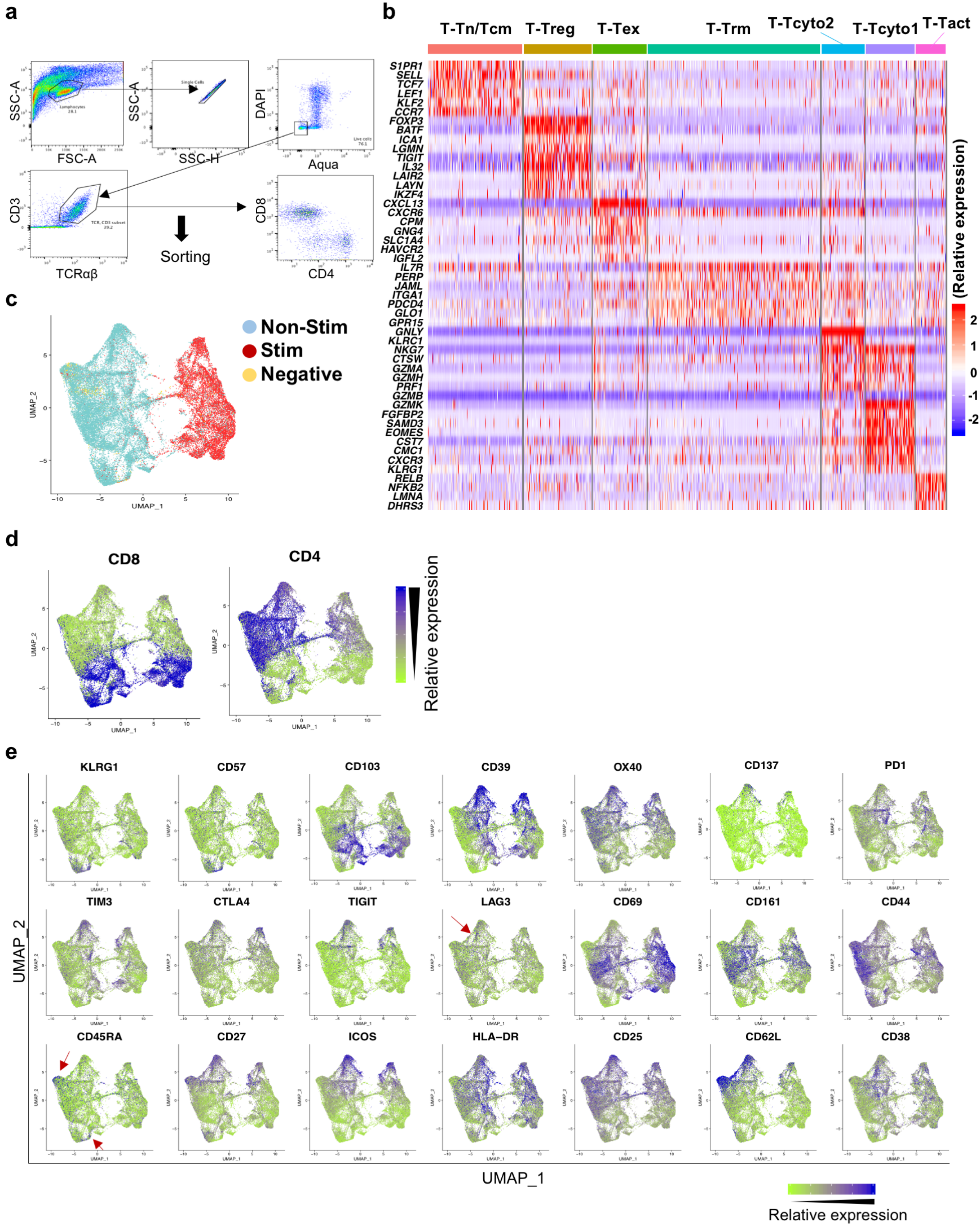


Extended Data Fig.1

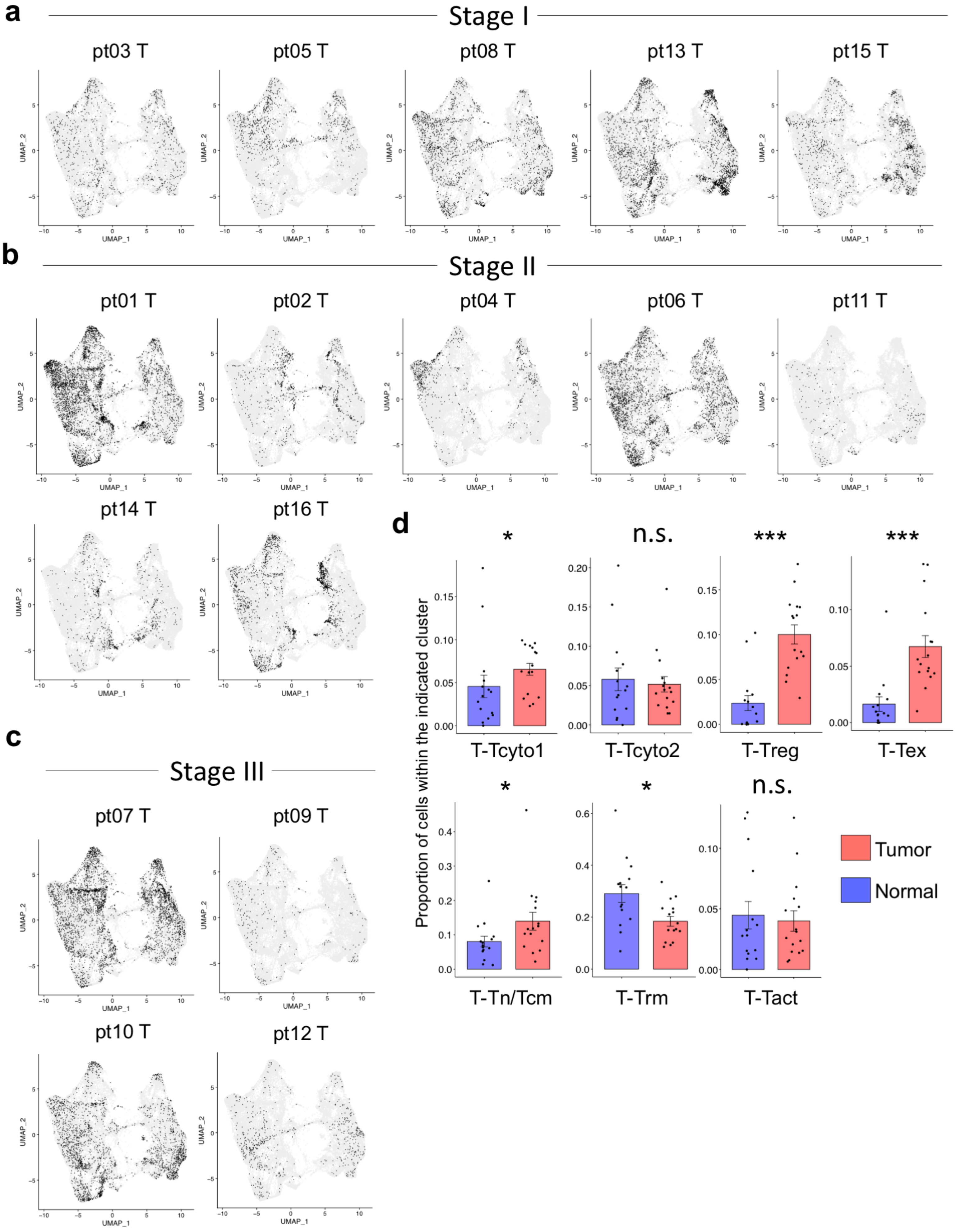


Extended Data Fig.2

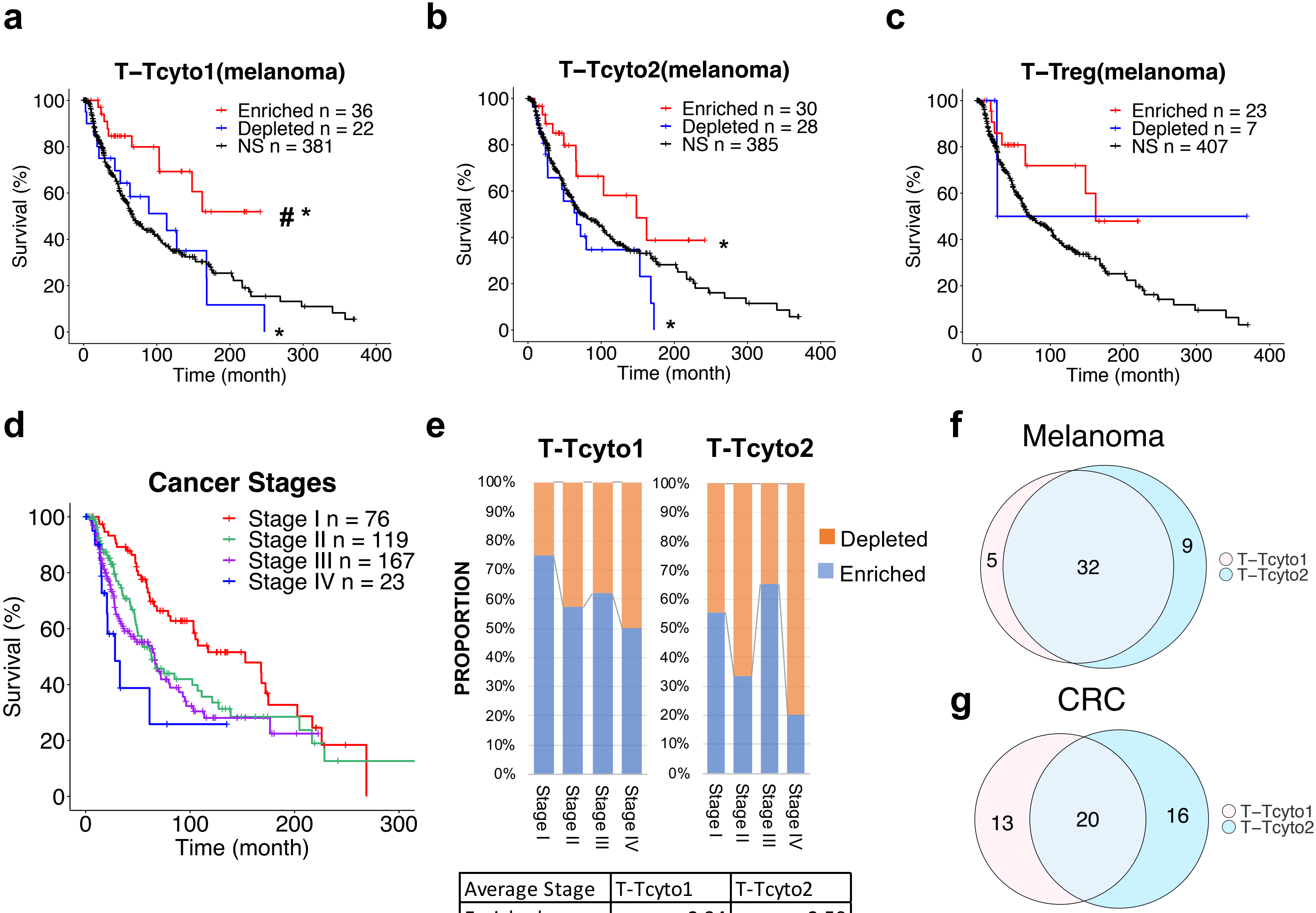


| Average Stage | T-Tcyto1 | T-Tcyto2 |
|---|---|---|
| Enriched | 2.34 | 2.50 |
| Depleted | 2.56 | 2.55 |

Extended Data Fig.3

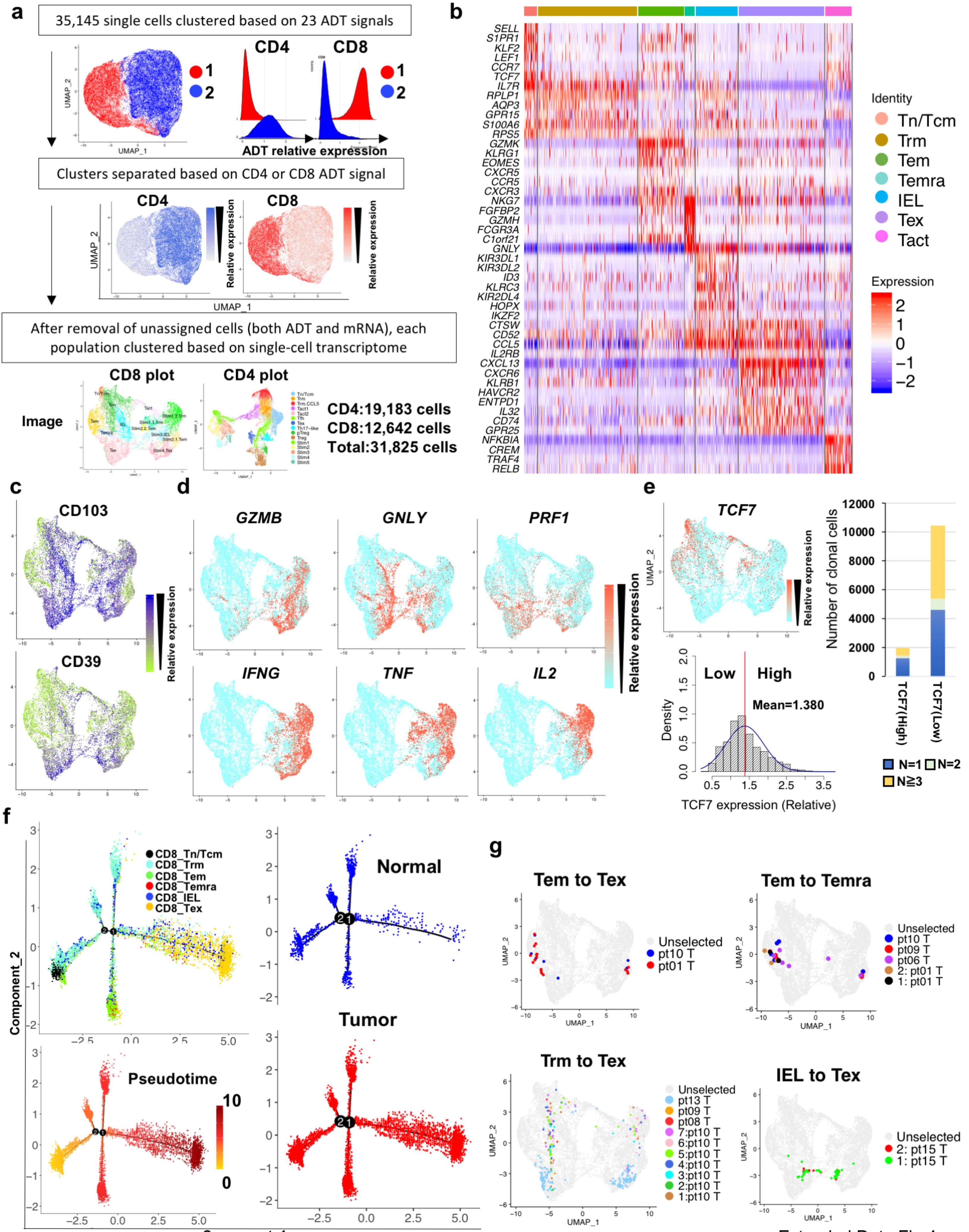


Extended Data Fig.4

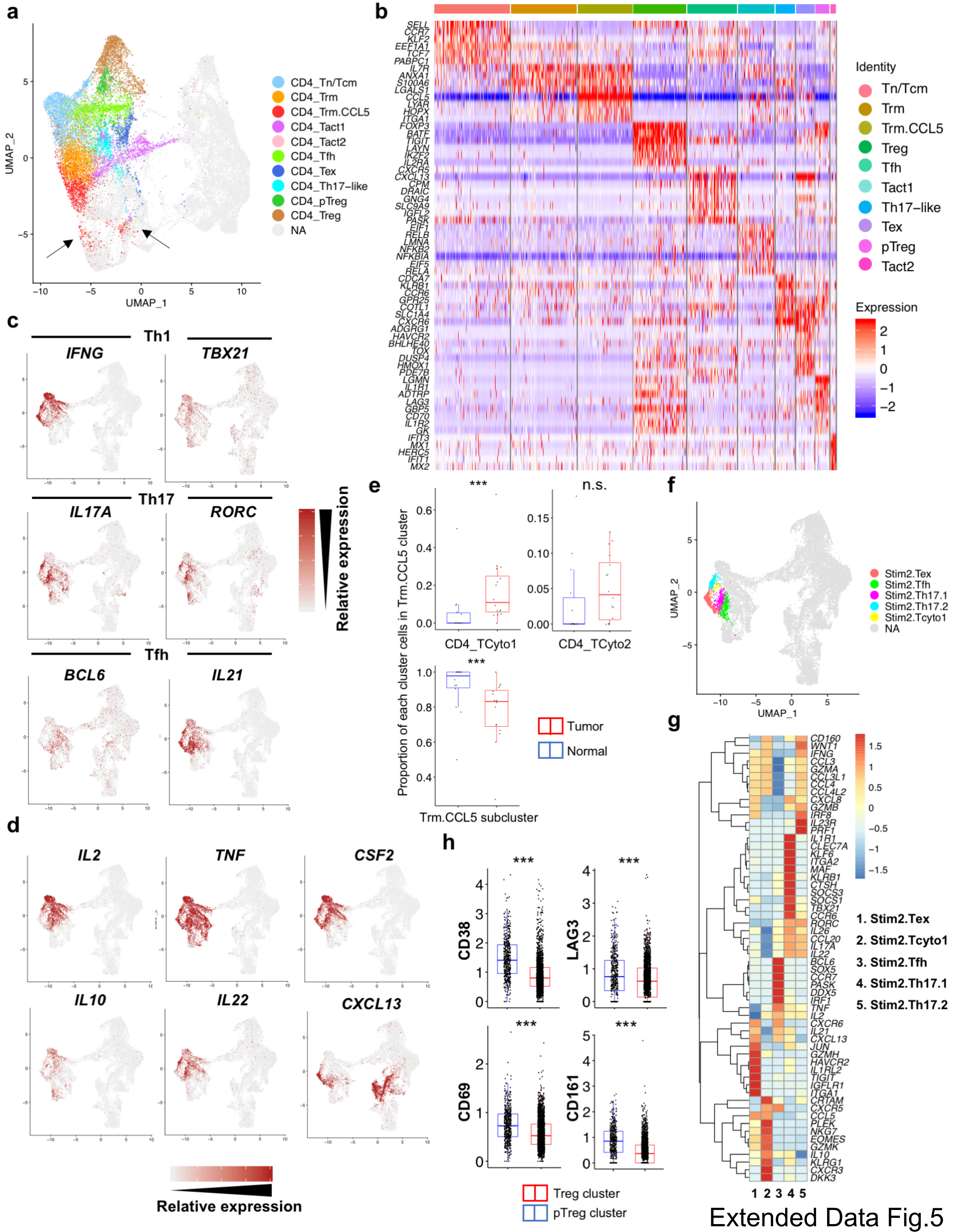


Extended Data Fig.5

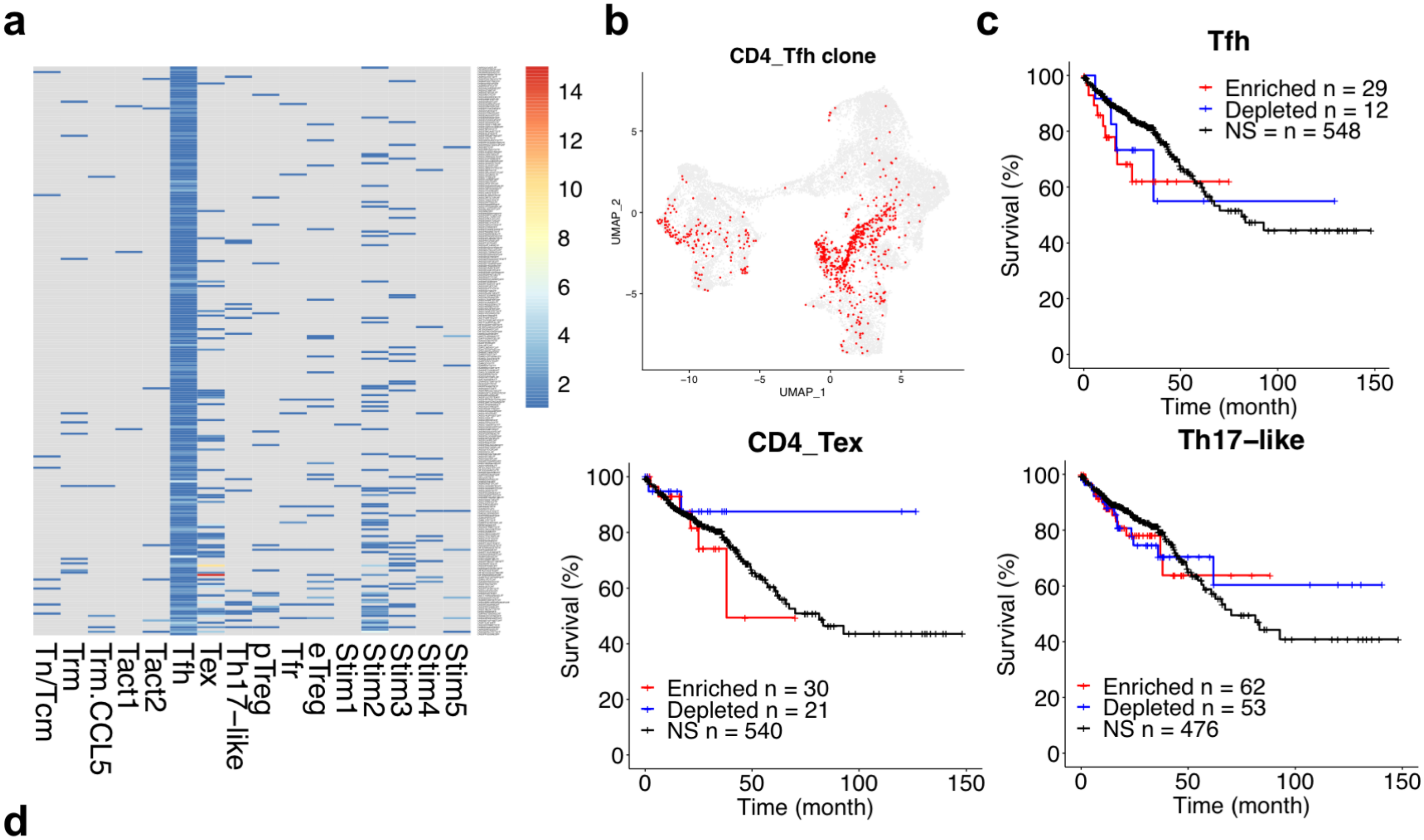


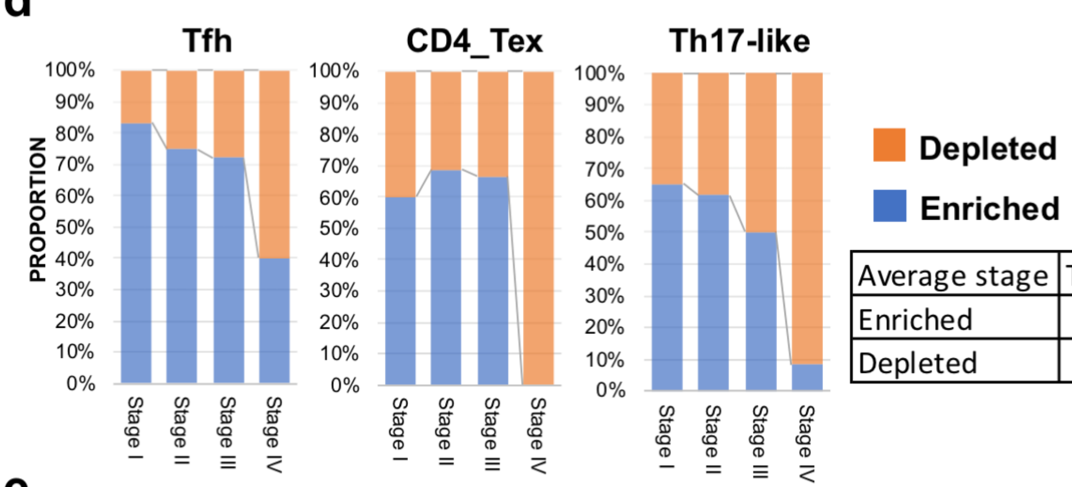


| Average stage | Tfh | CD4_Tex | Th17-like |
|---|---|---|---|
| Enriched | 2.41 | 2.03 | 2.05 |
| Depleted | 2.83 | 2.43 | 2.57 |

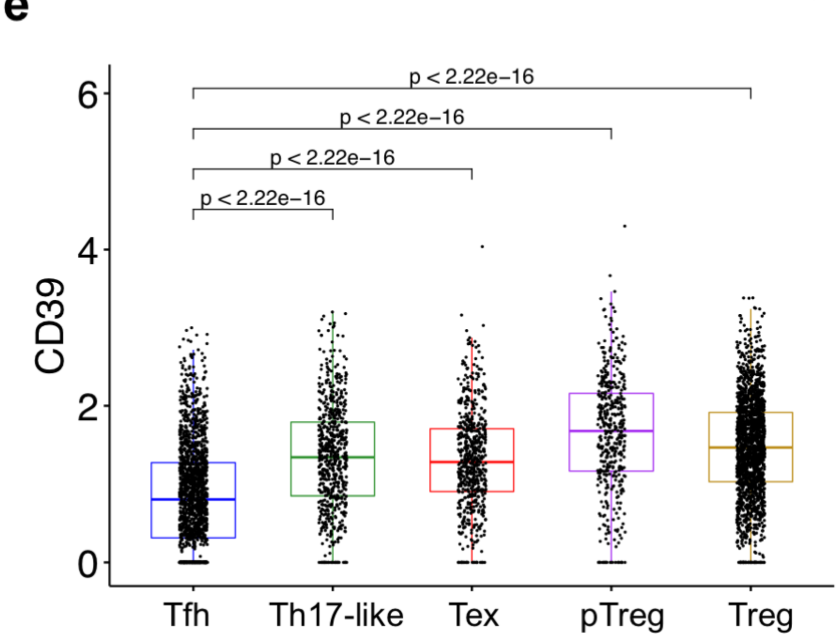


Extended Data Fig.6

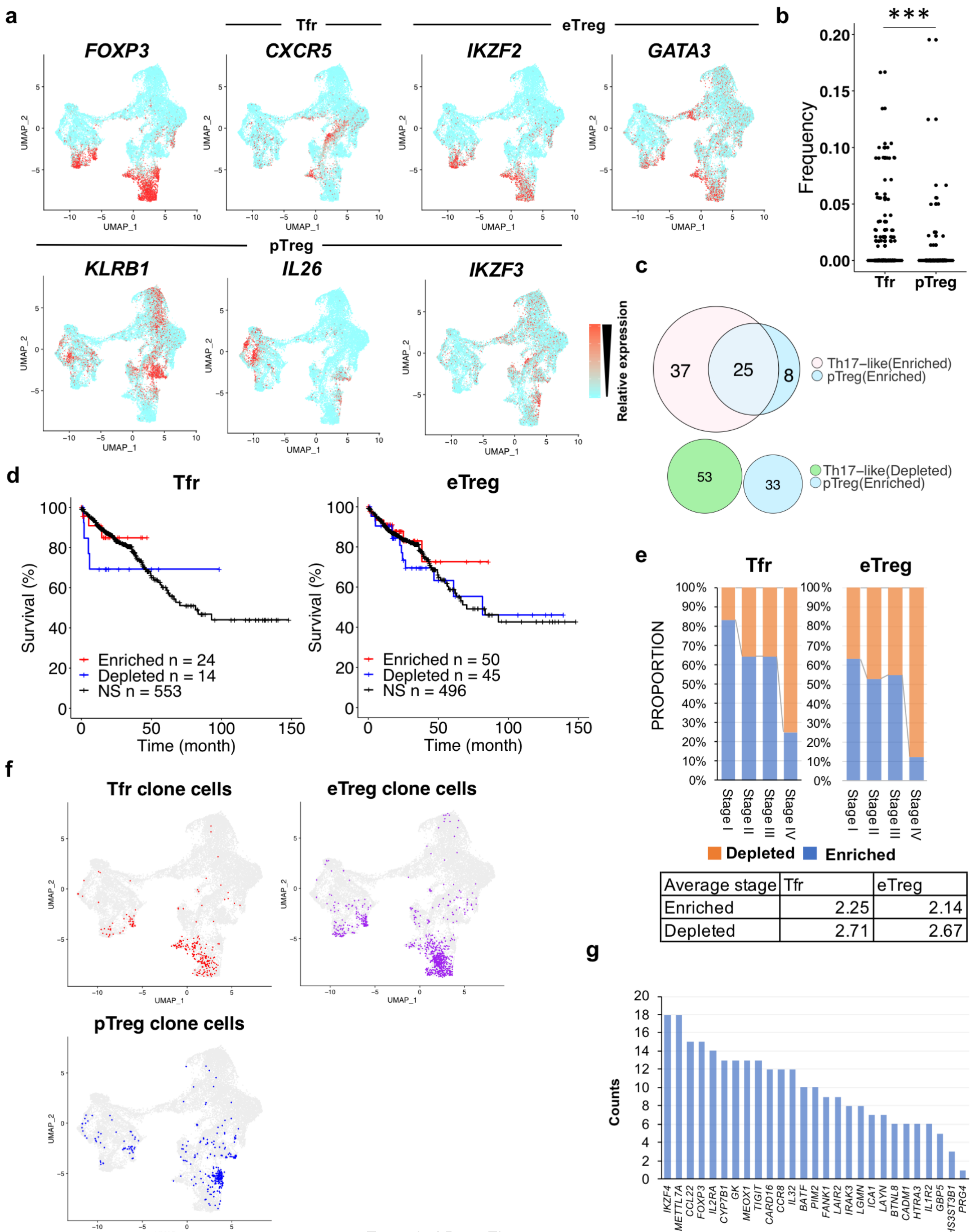


| Average stage | Tfr | eTreg |
|---|---|---|
| Enriched | 2.25 | 2.14 |
| Depleted | 2.71 | 2.67 |

Extended Data Fig.7
